# NAC1 Modulates Autoimmunity by Suppressing Regulatory T Cell-Mediated Tolerance

**DOI:** 10.1101/2022.03.01.482525

**Authors:** Jin-Ming Yang, Yijie Ren, Anil Kumar, Xiaofang Xiong, Jugal Kishore Das, Hao-Yun Peng, Liqing Wang, Xingcong Ren, Yi Zhang, Cheng Ji, Yan Cheng, Li Zhang, Robert C. Alaniz, Paul de Figueiredo, Deyu Fang, Xiaoqi Liu, Jianlong Wang, Jianxun Song

**Affiliations:** Department of Toxicology and Cancer Biology, Department of Pharmacology and Nutritional Science, and Markey Cancer Center, University of Kentucky College of Medicine, Lexington, KY 40536, USA; Department of Microbial Pathogenesis and Immunology, Texas A&M University Health Science Center, Bryan, TX 77807, USA; Department of Veterinary Pathobiology, Texas A&M University; Norman Borlaug Center, Texas A&M University, College Station, TX 77845, USA; Department of Pathology, Northwestern University Feinberg School of Medicine, Chicago, IL 60611, USA; Department of Medicine, Columbia Center for Human Development, Columbia University Irving Medical Center, New York, NY 10032, USA

## Abstract

FoxP3^+^ regulatory T cells (Tregs) are a distinct subset of CD4^+^ T cells integral to the maintenance of the balance of the immune system, and their dysregulation is a trigger of autoimmunity. We report here that nucleus accumbens-associated protein-1 (NAC1), a nuclear factor of the <u>b</u>road complex, tramtrack, <u>b</u>ric-a-brac / <u>po</u>xvirus and <u>z</u>inc finger (BTB/POZ) gene family, is a negative regulator of FoxP3 in Tregs and a critical determinant of immune tolerance. Phenotypically, NAC1^-/-^ mice show substantial tolerance to the induction of autoimmunity, as evidenced by the significantly decreased occurrences of autoimmune arthritis and colitis. Analysis of T cells from the wild-type (WT) or NAC1 knockout (^-/-^) mice found that NAC1 is crucially involved in the early stage of T cell development. NAC1 positively affects CD8^+^ T cell differentiation, but negatively regulates Treg development. Compared with WT animals, NAC1^-/-^ mice displayed defects in CD8^+^ T cell development but generated a larger amount of CD4^+^ regulatory Tregs that exhibit a higher metabolic profile and immune suppressive activity, increased acetylation, and expression of FoxP3, and slower turnover of this transcriptional factor. Furthermore, treatment of Tregs with the pro-inflammatory cytokines IL-1*β* or TNF-*α* induced a robust upregulation of NAC1 but an evident downregulation of FoxP3 as well as the acetylated FoxP3, suggesting that the reduction of FoxP3 by the NAC1-mediated deacetylation and destabilization of this lineage-specific transcriptional factor contributes considerably to break of immune tolerance. These findings imply that the pro-inflammatory cytokines-stimulated upregulation of NAC1 acts as a trigger of the immune response through destabilization of Tregs and suppression of tolerance induction, and that therapeutic targeting of NAC1 warrants further exploration as a potential tolerogenic strategy for treatment of autoimmune disorders.

## Introduction

Aberrant autoimmunity results in over 80 different autoimmune diseases that are often debilitating and life-threatening, for which there is no cure at present. Autoimmune diseases such as Crohn’s disease, type-1 diabetes mellitus (T1D), rheumatoid arthritis (RA) and ulcerative colitis (UC) are believed to result from interaction between genetic and environmental factors and to be a consequence of compromised immune tolerance versus adaptive immune response. Immune tolerance prevents an immune response to a particular antigen (Ag) or tissues that cause autoimmune disorders, and a range of immune cell types participate in the control of hyposensitivity of the adaptive immune system to the self-Ag or non-self-Ag.

Among these immune cells, FoxP3^+^ regulatory T cells (Tregs), a distinct and dynamic subset of CD4^+^ T cells, are an essential contributor to the immune tolerance, maintenance of immune cell homeostasis and the balance of the immune system; and defects in Tregs occur in virtually all the autoimmune disorders (*1*). The stability of the suppressor Tregs is critical for their function but is reduced in most of the autoimmune disorders. Therefore, maintenance of the Treg stability is crucial for immunologic tolerance. Yet, how impaired balance between immune response and tolerance is triggered and the key molecular determinants that affect Treg stability remain elusive.

Here, we report our new finding that nucleus <u>ac</u>cumbens-associated protein-1 (NAC1), encoded by *NACC1* gene and originally identified as a cocaine-inducible transcript from the nucleus accumbens (*2*), acts as a vital modulator of immune suppression. NAC1 is a nuclear factor that belongs to the BTB (Broad-Complex, Tramtrack and Bric a brac)/POZ (POX virus and Zinc finger) gene family. Using NAC1-deficient (^-/-^) mice, we uncovered a previously unrecognized but important role of NAC1 in triggering autoimmunity and Treg instability and demonstrated that NAC1 contributes to break of immune tolerance through its negative control of Treg development and function associated with deacetylation and destabilization of FoxP3 protein.

## Results

### Overall T cell population is divergent in the WT and NAC1^-/-^ mice

Previous studies have shown that NAC1 participates in regulation of the self-renewal and pluripotency of embryonic stem cells (*3–5*) and somatic cell reprogramming (*6*), and we recently found that NAC1 has a critical role in cellular metabolism (*7*). As metabolic reprogramming can significantly influence T cell activation, expansion, and effector function (*8, 9*), we queried whether NAC1 affects T cell development and function. We first performed T cell profiling in WT and NAC1^-/-^ mice. Compared with WT mice, development of T cells in the thymus of NAC1^-/-^ mice was curbed, as evidenced by increased numbers of thymocytes in the dominant-negative (DN) stage (1.67% *vs*. 0.72%) and decreased numbers of cells in the DN4 stage (4.65% *vs*. 28.5%; *p*<0.0001) (**Fig. 1A**). Although NAC1^-/-^ mice showed a reduction of total thymocytes and decreased cell amount in the DN4 stage, an accumulation of T cells in the DN2 stage was observed in those animals (49.1.5% *vs*. 28.7%; *p*<0.0001) (**Fig. 1A, Fig. 1B**). These alterations were correlated with a decreased percentage of TCRβ^+^ cells in DN4 cells found in NAC1^-/-^ mice. Despite a higher % of DN cells in NAC1^-/-^ mice, which show higher numbers of CD117^+^ cells, WT animals had a greater number of TCR*β*^+^ cells than NAC1^-/-^ mice. Like the gating on the DN4 population, there was a decrease in TCR*β*^+^ cells in NAC1^-/-^ mice as compared with WT animals, which could be due to a higher % of DN4 cells in WT mice (**Extended Fig. 1**). Conversely, in the lymph nodes (LNs) and spleen of NAC1^-/-^ mice, there was a reduced percentage of CD8^+^ T cells (28.5% *vs.* 40.5%) but an increased percentage of CD4^+^ T cells (66.3% *vs.* 57.3%) as compared to the controls (**Fig. 1C**). Despite the similar numbers of the total CD4^+^ or CD8^+^ single positive (SP) T cells in the thymus (**Fig. 1B**; lower panel), there were significant increases of % and numbers of CD4^+^ SP T cells (*p*<0.05 or *p*<0.01) but a decrease of CD8^+^ SP T cells (*p*<0.05 or *p*<0.01) in the pooled LNs and spleen in NAC1^-/-^ mice, as compared with WT controls (**Fig. 1D; lower panel**). These results suggest an important role for NAC1 in the early stage of T cell development and in the differentiation of CD4^+^ and CD8^+^ SP cells.

**Figure 1.**
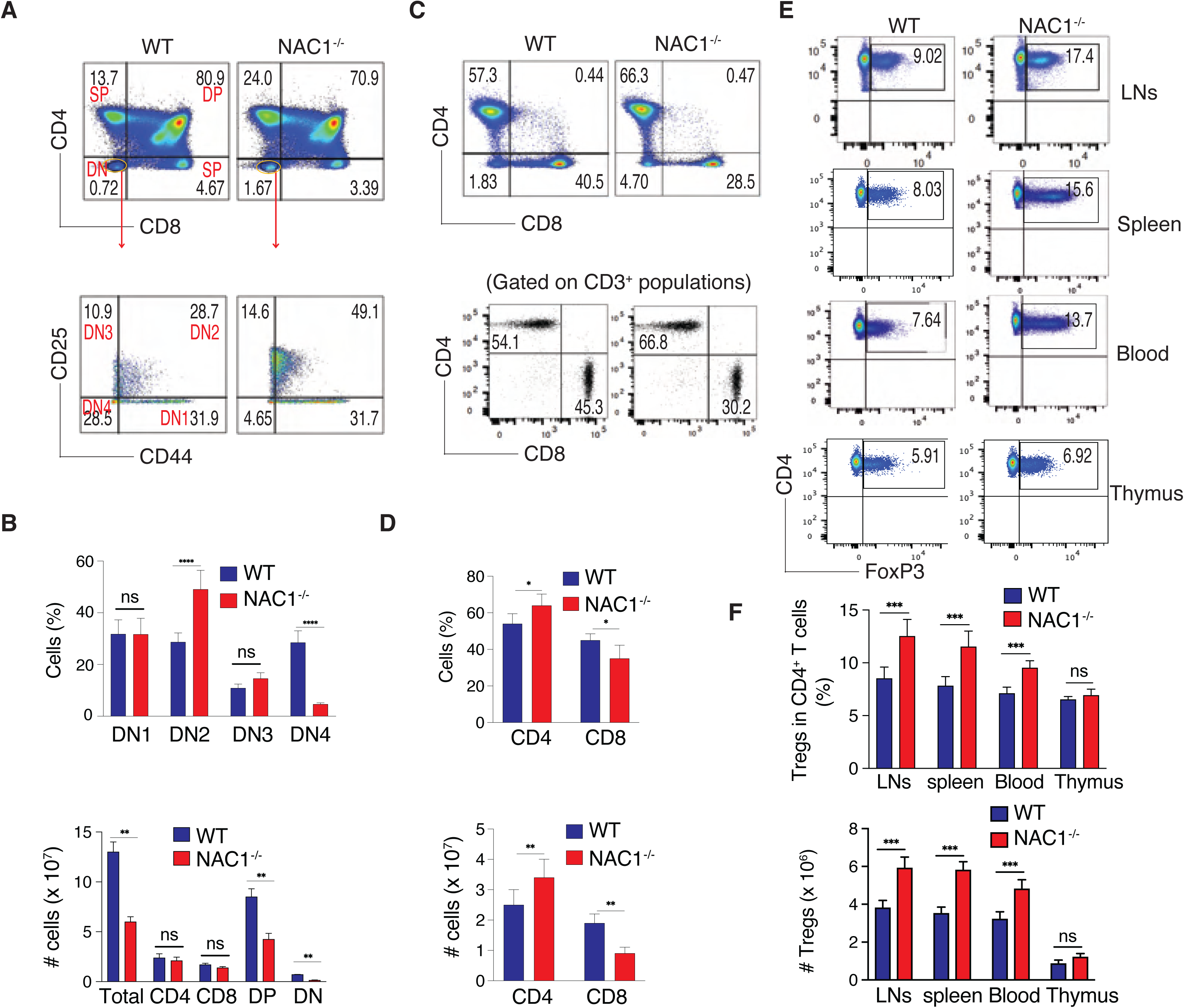
Loss of NAC1 impacts overall T cell populations. T cells from the thymus, peripheral lymph nodes (LNs) and spleen of WT or NAC1^-/-^ mice were analyzed by flow cytometry and calculated for numbers or percentages. (**A**) CD4 and CD8 in the thymus. The double negative (DN) populations were analyzed for DN1 to DN4 stages based on CD44 and CD25. Data shown are the representative of five mice per group of three independent experiments. (**B**) Numbers and percentages of total thymocytes, CD4 or CD8 single positive (SP) and the percentages of DN2 or DN4 cells. Data shown are the representative of three identical experiments. The values represent mean ± S.D. (N = 4 or 5). ****, *p*<0.0001, ns, no difference, Student’s unpaired *t*-test. (**C**) CD4 and CD8 T cells from the pooled LNs and spleen. Data shown are the representative of five mice per group of three independent experiments. ***, *p*<0.001, Student’s unpaired *t*-test. (**D**) Numbers and percentages of T cells from the pooled LNs and spleen. Data shown are representative of three identical experiments. The values represent mean ± S.D. (N = 4 or 5). *, *p*<0.05, **, *p*<0.01, Student’s unpaired t-*t*est. (**E**) Representative CD4^+^FoxP3^+^ Tregs in the pooled LNs, spleen, blood, and thymus, gating on CD4^+^ populations. Data shown are the representative of three identical experiments (N = 4 or 5). (**F**) Numbers and percentages of Tregs. Data shown are the representative of three identical experiments. The values represent the mean ± S.D. (N = 4 or 5). ***, *p*<0.001; ns, no statistical difference, Student’s unpaired *t*-test.

### Deficiency of NAC1 promotes Treg development and stability

The significant increase of total peripheral CD4^+^ T cell population observed in NAC1^-/-^ mice (**Fig. 1D**) prompted us to ask whether NAC1 plays a regulatory role in the development of Tregs, a unique sub-type of CD4^+^ T cells able to suppress excessive immune reaction. Indeed, as compared with WT animals, NAC1^-/-^ mice showed an evident increase in % of Tregs in the LNs, spleen and blood (**Fig. 1E**). Moreover, NAC1^-/-^ animals had significantly higher % and numbers of Tregs in the LNs and spleen but not in the thymus (*p*<0.0001; **Fig. 1F**). To prove the role of NAC1 in the development of Tregs, we used an *in vitro* system in which induced Tregs (iTregs) are generated from naive CD4^+^CD25^-^ T cells (*10*). The naive CD4^+^CD25^-^ T cells from the LNs and spleen of WT or NAC1^-/-^ mice were treated with TGF-*β* to produce iTregs. The naive CD4^+^CD25^-^ T cells from WT mice expressed abundant NAC1 but no detectable FoxP3; notably, the iTregs from those T cells showed a robust expression of FoxP3 but a substantial reduction of NAC1 expression (**Fig. 2A**). Remarkably, a significantly greater amount of iTregs were generated from NAC1^-/-^ than from WT CD4^+^CD25^-^ T cells (**Fig. 2B-C**). Furthermore, in response to stress, the expression of CD36, an immuno-metabolic receptor that mediates metabolic adaptation and supports Treg survival and function (*11, 12*), was significantly elevated in NAC1^-/-^ Tregs as compared with WT Tregs (**Fig. 2D-E**). These results indicate that NAC1 exerts a negative control of Treg development at early stages. In addition to affecting CD4^+^ populations, NAC1 also affects CD8^+^ T cells. NAC1 deficiency led to a significant decrease of CD8^+^ SP T cell generation in the LNs and spleen, reduced production of cytokine, and shortened cellular survival (**Extended Fig. 2**). Viral Ag-stimulated development of memory CD8^+^ T cells was also suppressed in the absence of NAC1 (**Extended Fig. 3**).

**Figure 2.**
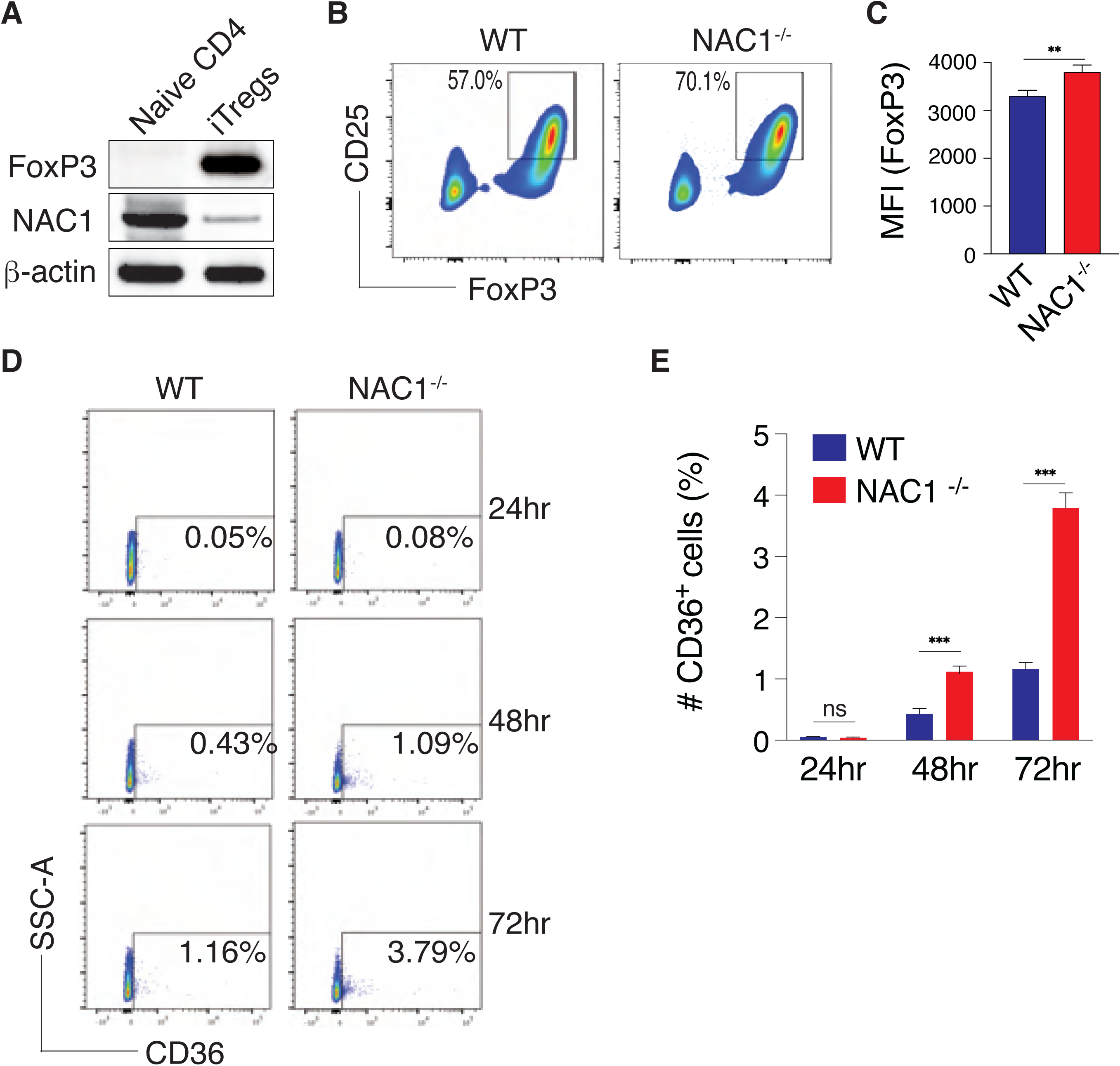
Loss of NAC1 enhances the induction of iTregs and expression of CD36. The naive CD4^+^CD25^-^ T cells from the pooled spleen and LNs of WT or NAC1^-/-^ mice were induced to iTregs *in vitro* in the presence of TGF-*β* for 5 days. (**A**) Expression of NAC1 in naive CD4 and iTregs of WT T cells detected by immunoblots. (**B**) Expression of CD25 and FoxP3 in iTregs generated from WT and NAC1^-/-^ CD4^+^CD25^-^ T cells by flow cytometry. Data shown are the representative of three identical experiments. (**C**) Mean Fluorescence Intensity (MFI) of FoxP3 in iTregs generated from WT and NAC1^-/-^ CD4^+^CD25^-^ T cells as analyzed by flow cytometry. Results shown are the mean ± S.D. of three identical experiments. **, *p*<0.01, Student’s unpaired *t*-test. (**D**) CD36 expression of WT or NAC1^-/-^ iTregs following lactic acid treatment. Tregs generated *in vitro* were treated with 10mM lactic acid for various times and analyzed by flow cytometry for CD36 expression. Data shown are the representative of three identical experiments. (**E**) Quantification of percentage of CD36^+^ Tregs following lactic acid treatment. Results shown are the mean ± S.D. of three identical experiments. *** *P*<0.001, Student’s unpaired *t*-test.

### NAC1^-/-^ Tregs display enhanced functional activities

To further validate the negative control of Treg development by NAC1, we examined the functional activity of the Tregs either from WT or NAC1^-/-^ mice. CD4^+^CD25^+^ Tregs from the LNs and spleen of WT or NAC1^-/-^ mice were stimulated with the mouse CD3/CD28-loaded beads in the presence of rIL-2, and the metabolic differences between WT or NAC1^-/-^ cells were then analyzed using the Seahorse XF Cell Mito Stress Test kit. Although Tregs from WT or NAC1^-/-^ mice had similar proliferation and survival profiles (**Fig. 3A-C**), Tregs from NAC1^-/-^ mice exhibited a significantly higher oxygen consumption rate (OCR) and glycolytic rate (glycoPER) than Tregs from WT mice (**Fig. 3D-E**), suggesting that NAC1^-/-^ Tregs are metabolically more active than the corresponding control Tregs. Consistently, NAC1^-/-^ Tregs produced significantly greater amounts of the suppressive cytokines, TGF-*β* and IL-10, than WT Tregs, as shown by intracellular staining (**Fig. 3F)** and ELISA (**Fig. 3G**), which may constitute one of several mechanisms that contribute to the suppressive capacity of Tregs. These results clearly demonstrate a negative impact of NAC1 on the suppressive function of Tregs and imply that inhibiting NAC1 may modulate autoimmunity through promoting Treg development and function. The enhanced function of NAC1^-/-^ Tregs was further demonstrated in an *in vitro* suppressive assay (**Fig. 3H**) and in an autoimmune colitis model subjected to *in vivo* co-transfer of Tregs with CD4^+^ T effectors (Teffs), which showed that NAC1^-/-^ Tregs elicited a stronger immune suppressive effect on inflammation than WT Tregs. Transfer of naive Teffs into *Rag*1^-/-^ mice resulted in weight loss in two weeks, and co-transfer of Teffs with FoxP3^+^ Tregs (Treg/Teffs ratio of 1:3) from NAC1^-/-^ mice led to a significant reduction of weight loss compared with those of WT mice (*p*<0.05; **Fig. 3I**). The co-transfer of Teffs with NAC1^-/-^ and WT FoxP3^+^ Tregs effectively protected from a decrease in finger-like villus projections (arrows) in the gut (**Fig. 3J**). Similarly, in the LPS-induced colitis, adoptive transfer of NAC1^-/-^ Tregs was more efficient in preventing colon loss than WT Tregs, as shown by the colon lengths (**Fig. 3K-L**), histology (an arrow indicates the goblet cell architecture of the colon tissue was damaged by LPS; **Fig. 3M**) and Treg infiltration in the draining LNs (**Fig. 3N**). Furthermore, we performed *in vivo* analyses on survival, proliferation/cell cycle, and apoptosis. WT and NAC1^-/-^ Tregs (Thy1.2^+^) were labeled with CFSE and *i.v.* injected into B6.Thy1.1 Tg mice. On Days 2 and 5 after Treg transfer, the pooled LNs and spleen were analyzed for the transferred Th1.2^+^ Tregs using flow cytometry. We found that NAC1^-/-^ Tregs had similar survival and proliferation/cell cycle (**Fig. 3O**), but increased FoxP3 expression (**Fig. 3P)** and reduced apoptosis (**Fig. 3Q**) as compared with WT Tregs. In addition, we found that both NAC1^-/-^ iTregs and nTreg produced more suppressive cytokines (i.e., IL-10 and TGF-*β*) than WT cells, and the similarity between iTregs and nTregs (**Extended Fig. 4**). In a mouse tumor model receiving the *in vivo* co-transfer of Tregs with CD8^+^ T cells, NAC1^-/-^ Tregs displayed greater suppressive effect on antitumor immunity than the control Tregs (**Extended Fig. 5**), another evidence for the enhanced suppressive activity of NAC1^-/-^ Tregs.

**Figure 3.**
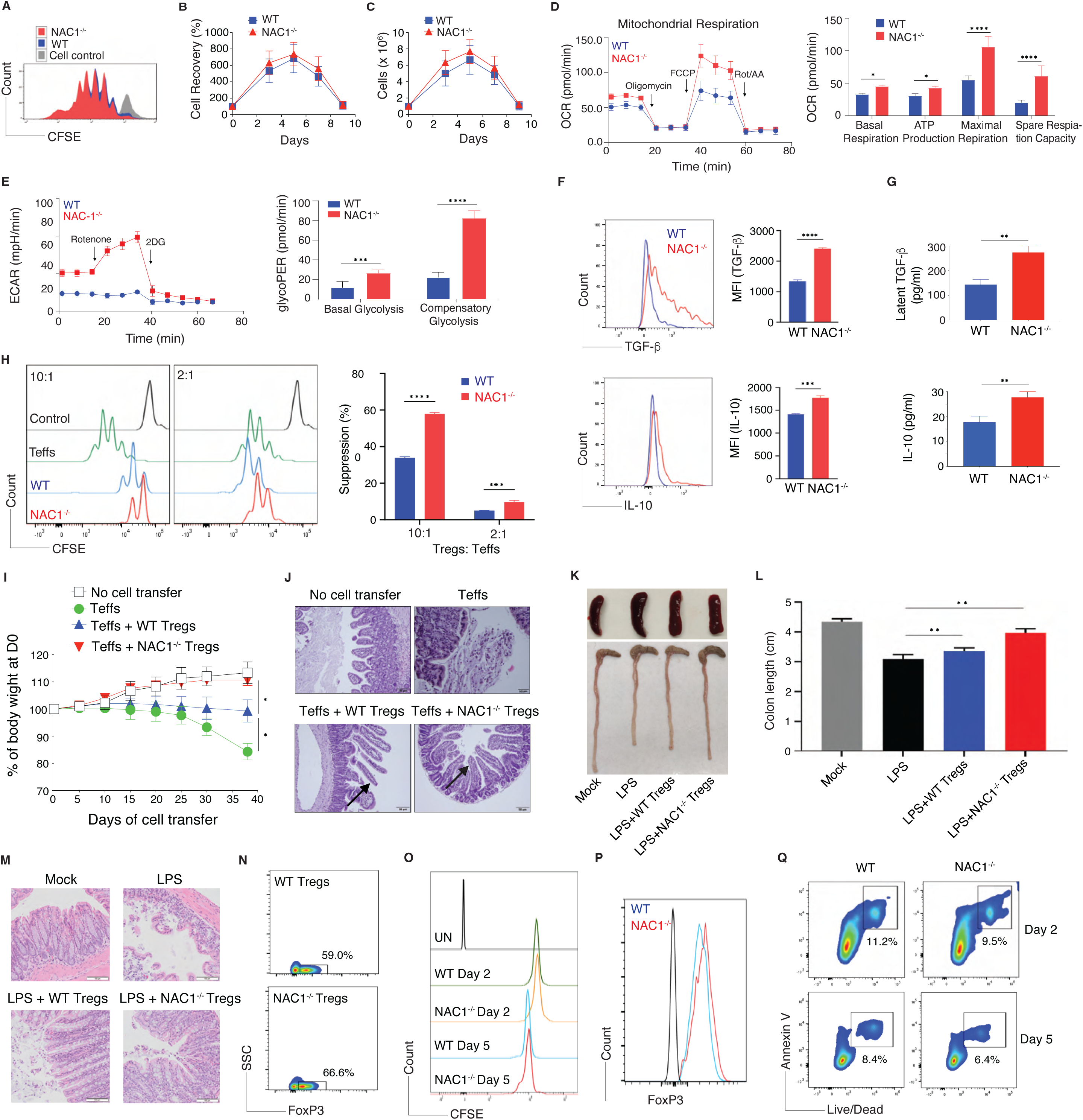
Loss of NAC1 enhances the functional activity of Tregs. Purified CD4^+^ Tregs from the pooled LNs and spleen of WT or NAC1^-/-^ mice were stimulated with anti-CD3 plus CD28 antibodies in the presence of rIL-2 for various time. (**A**) Cell proliferation/division by CFSE-based flow cytometry. Data shown are the representative of three identical experiments. (**B**) % and (**C**) numbers of cell recovery on various days were examined by trypan blue exclusion. The numbers of T cells present on day 0 were assigned a value of 100%, and numbers surviving on various days were used to calculate the percentage recovery relative to day 0. Data shown represent the mean ±.E.M. of percentage change or numbers of three independent experiments. All *P*>0.05, Student’s unpaired *t*-test). (**D**) Oxygen consumption rate (OCR, y axis) trace and tabulated data (right) of Tregs. * *P*<0.05, ****, *P*<0.0001, Student’s unpaired *t*-test. (**E**) Glycolytic rate (glycoPER, y axis) trace and tabulated data (right) of Tregs. *** *P*<0.01, ****, *P*<0.0001, Student’s unpaired *t*-test. (**F**) Cytokine production of 16 hr by intracellular staining. Data shown are the mean ± S.D. of three independent experiments (N = 3). *** *P*<0.001; **** *P*<0.0001, Student’s unpaired *t*-test. (**G**) Cytokine secretion of 48 hr by ELISA. Data shown are the mean ± S.D. of three independent experiments (N = 3). ** *P*<0.01, Student’s unpaired *t*-test. (**H**) *In vitro* suppressive assay. CD4^+^ CD25^-/-^ Teffs pre-labelled with CFSE were non stimulated (control) or stimulated with anti-CD3 plus CD28 antibodies in the absence of WT or NAC1^-/-^ Tregs (10:1 or 2:1) for three days. Cell proliferation was analyzed by flow cytometry, gating on CFSE^+^ population (left panel) and % suppression was calculated (right panel). Data shown are the mean ± S.D. of three independent experiments (N = 3). *** *P*<0.001; **** *P*<0.0001, Student’s unpaired *t*-test. (**I-J**) T cell transfer model of colitis. (**I**) Changes of body weight of *Rag*1^-/-^ host mice after adoptive cell transfer of naive CD4^+^ Teffs with or without Tregs from WT or NAC1^-/-^ mice. Data shown are the mean ± S.E.M. of body weight change from a representative of three identical experiments (N = 10). *, *p*<0.05, Nested *t*-test). (**J**) Representative H&E-stained sections of gut tissues collected from the recipient mice 4 weeks after adoptive cell transfer. (**K-N**) LPS-mediated colitis. (**K**) Representative colon images of the LPS-induced colitis from two identical experiments (N = 5). (**L**) Colon lengths in LPS-induced colitis. Data shown are the representative of two identical experiments (N = 5). The values represent the mean ± S.D. ** *P*<0.01, Student’s unpaired *t*-test. (**M**) Representative H&E-stained sections of images of proximal colon cross-section of mock or LPS-challenged B6.Thy1.1 Tg mice with transferred WT or NAC1^-/-^ Tregs at day 8 of two identical experiments (N = 5). (**N**) Flow cytometric analysis of transferred FoxP3^+^ Tregs in draining LNs, gating on Thy1.2^+^ populations. Data shown are the representative of two identical experiments (N = 5). (**O-Q**) *In vivo* analyses of Tregs. WT and NAC1^-/-^ Tregs (Thy1.2^+^) were labelled with CFSE and *i.v.* injected into B6.Thy1.1 Tg mice. On Days 2 and 5 after Treg transfer, the pooled LNs and spleen were analyzed for the transferred Th1.2^+^ Tregs by flow cytometry. (**O**) Proliferation by CFSE. Data shown are the representative of two identical experiments (N = 5). (**P**) FoxP3 expression. Data shown are representative of two identical experiments (N = 5). (**C**) Apoptosis analysis by Annexin V and PI. Data shown are the representative of two identical experiments (N = 5).

### NAC1^-/-^ mice are insusceptible to induction of autoimmunity

To further prove the impact of NAC1 on autoimmunity, next we compared the response of the WT and NAC1^-/-^ mice to induction of autoimmune arthritis and colitis. Type II collagen was used to induce arthritis (*13*) and dextran sulfate sodium (DSS) was given to mice to induce colitis (*14*). We found that NAC1^-/-^ mice were significantly tolerant to induction of autoimmune arthritis and colitis (**Fig. 4**). In the collagen-induced arthritis model (CIA) (*13*), a significantly lower occurrence of CIA was observed in NAC1^-/-^ mice than in the littermate controls, as determined by the histologic evidence (**Fig. 4A**), disease incidence (**Fig. 4B;** *p*<0.0001) and disease score (**Fig. 4C;** *p*<0.0001). Tolerance to autoimmunity induction was recapitulated in a colitis model in which mice were given drinking water containing dextran sulfate sodium (DSS). WT mice developed autoimmune colitis within 3-4 days following DSS administration and showed visible signs of illness including hunched back, raised fur, symptoms of sepsis and reduced mobility because of diarrhea and anemia; strikingly, the occurrence of colitis declined remarkably in NAC1^-/-^ mice (**Fig. 4D-H**). In NAC1^-/-^ mice, the body weight loss (**Fig. 4E**), survival (**Fig. 4F**), colon shrinkage (**Fig. 4G**) and the disease activity index (**Fig. 4H**) were all significantly improved as compared with WT animals (*p*<0.0001). In both of the disease models, there were considerably larger amounts of pro-inflammatory immune cells in the joint (**Fig. 4A**) and the colon (**Fig. 4D**) tissues of WT mice than in those of NAC1^-/-^ animals (arrows), suggesting that weakened immune response may account for the insusceptibility of NAC1^-/-^ mice to induction of autoimmunity. As NAC1^-/-^ mice showed enhanced numbers and functions of Tregs (**Fig. 1, Fig. 3**) and Tregs have a unique capacity to suppress immune response, the tolerance to the induction of autoimmune diseases observed in NAC1^-/-^ mice (**Fig. 4**) could be a consequence of the enhanced Treg stability.

**Figure 4.**
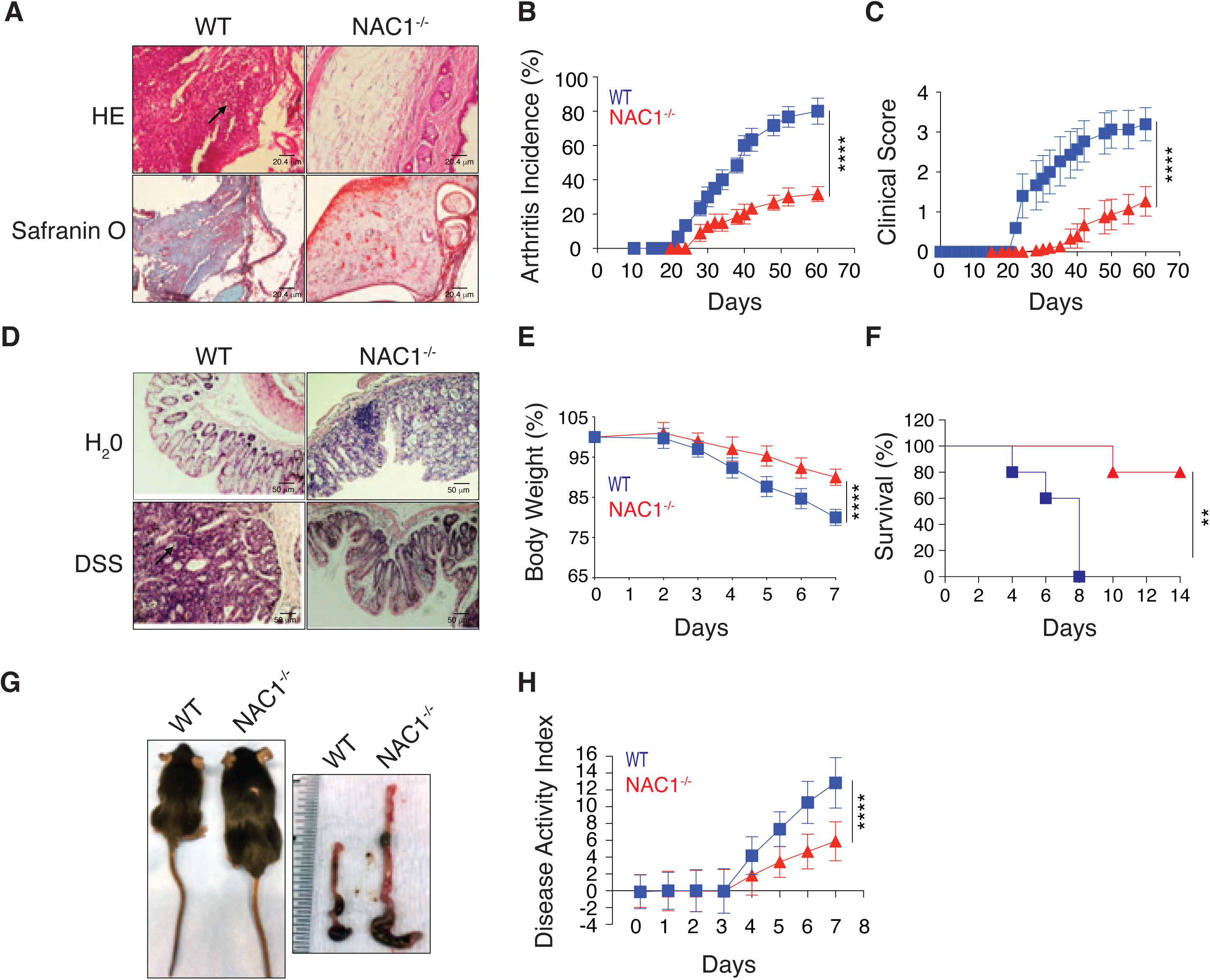
NAC1^-/-^ mice are tolerant to induction of autoimmunity. WT or NAC1^-/-^ mice were challenged with either bovine type II collagen in complete Freund’s adjuvant by one intradermal immunization at two sites in the base and slightly above of the tail on day 0, or by oral ingestion of 3% dextran sulfate sodium (DSS, MP Biomedicals) in drinking water for 5 days. (**A-C**) Arthritis: The histology of the joints (**A**), arthritis incidence (**B**), and clinical score (**C**) were evaluated by examining the paws. Values are the mean ± S.E.M. of three independent experiments (n=10). *P*<0.0001 in **B** and **C**, simple linear regression. (**D-H**) Colitis: The severity of colitis activity was graded on the designated dates. Histology of colon (**D**), animal body weight change (**E**), survival (**F**), animal size and colon length (**G**), and the resultant IBD disease activity index (**H**) were determined. Values are the mean ± S.E.M. of three independent experiments (n=10). *P*<0.0001 in E and H, simple linear regression. *P*<0.001 in F, survival curve comparison.

### DNA methylation of FoxP3 in Tregs remains intact in the absence of NAC1

Next, we sought to understand how NAC1 regulates the development and function of Tregs. As FoxP3 is a transcription factor essential for establishment and maintenance of Treg phenotype, and NAC1 may cooperate with certain transcriptional factors to regulate various cellular activity (*15, 16*), we therefore asked whether the effect of NAC1 on Tregs is mediated through its interaction with FoxP3.

Primarily, FoxP3 stability is associated with selective demethylation of an evolutionarily conserved element within the *FoxP3* locus named TSDR (Treg-specific demethylated region) (*17, 18*). To determine whether the effect of NAC1 on Tregs is mediated through FoxP3 methylation, we performed an analysis of TSDR demethylation. A panel of the FoxP3 loci (ADS779, ADS657, ADS569, ADS442, ADS443, ADS1183, ADS1184) between WT and NAC1^-/-^ Tregs were analyzed by the Targeted NextGen Bisulfite Sequencing (tNGBS) (**Fig. 5A**), and the CpG DNA methylation of FoxP3 in the four regions including Distal Region (ADS657, ADS569), Proximal Region (ADS1183), CNS2 Region (ADS443), and 3’ Downstream Region (ADS1184) were determined (**Fig. 5B**). No substantial differences of the CpG DNA methylation in the four regions were found between WT and NAC1^-/-^ Tregs (**Fig. 5C**, and **Extended Table 1**). The two bar graphs representing the specific methylation percentages of each FoxP3 CpG in WT and NAC1^-/-^ Tregs are comparably presented (**Fig. 5D**). These results indicate that NAC1 does not have detectable effects on epigenetic imprinting in the TSDR region of FoxP3.

**Figure 5.**
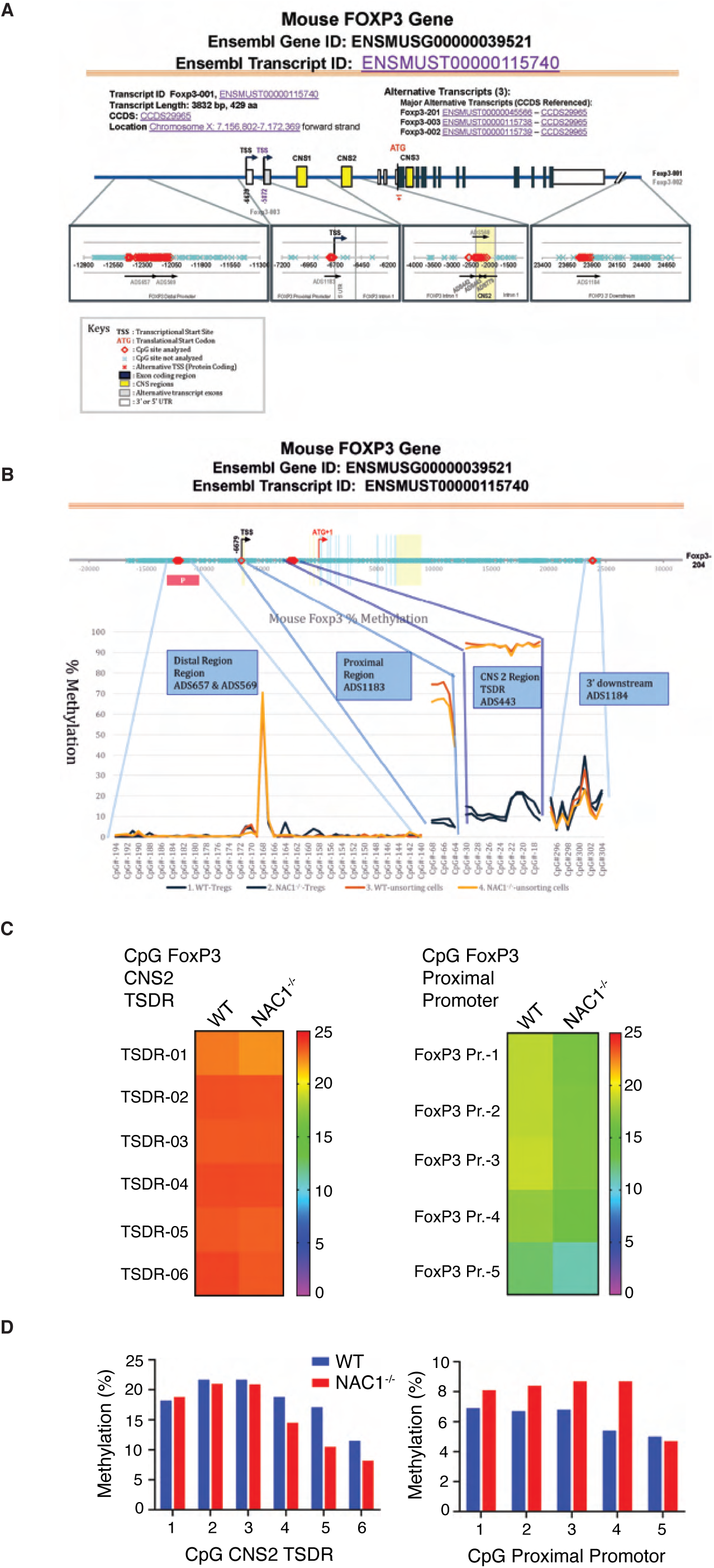
CpG motif analyses indicate that FoxP3 DNA methylation is not regulated by NAC1. Naive WT and NAC1^-/-^ CD4^+^CD25^+^Tregs were purified and used in these analyses. (**A**) Diagram of mouse FoxP3 DNA methylation. Seven CpG sites of FoxP3 regulators (ADS657, ADS569, ADS1183, ADS442, ADS443, ADS779 and ADS1184) were analyzed. (**B**) Four CpG regions of FoxP3 regulators, including Distal Region (Regions ADS657 and ADS569), Proximal Region (ADS1183), CNS 2 Region (TSDR ADS443) and 3’ downstream (ADS1184), were analyzed. (**C**) Genomic DNA of the lymphocytes from LNs and spleen of WT or NAC1^-/-^ mice was analyzed for methylation status of CNS2 (TSDR) and FoxP3 proximal promoter, respectively. The degree of methylation at each CpG motif is depicted according to the color code. (**D**) % methylation of each FoxP3 CpG based on (**C**). Data shown are the representative of two identical experiments.

### FoxP3 transcription in Tregs is unaltered in the absence of NAC1

As NAC1 is a transcription co-regulator, we wanted to know whether NAC1 affects the transcription of FoxP3. To compare FoxP3 transcription in Tregs with or without NAC1, we first examined the possible enrichment and direct binding of NAC1 at FoxP3 genetic locus in WT Tregs, using the chromatin immunoprecipitation coupled with deep sequencing (ChIP-seq). NAC1-associated chromatins were pulled down from the CD4^+^CD25^+^ Tregs of C57BL/6 mice, followed by a high throughput sequencing. The results showed that peak density of NAC1 was not enriched within regulatory elements of FoxP3 (**Extended Fig. 6; Fig. 6Ai**). Assay for transposase accessible chromatin sequencing (ATAC-seq) identified a total of over 10000 differential ATAC-seq peaks in WT and NAC1^-/-^ Tregs, but no significantly differential accessibility was found at FoxP3 gene locus (**Fig. 6Aii**). FoxP3 ChIP-seq validated that FoxP3 enrichment was not affected by NAC1 expression (**Fig. 6Aiii**). Furthermore, the RNA-seq showed that among ∼200 differentially expressed genes, FoxP3 was not differentially expressed in WT and NAC1^-/-^ Tregs (**Fig. 6B**). These results suggest that the effect of NAC1 on FoxP3 expression is not mediated at transcription level.

**Figure 6.**
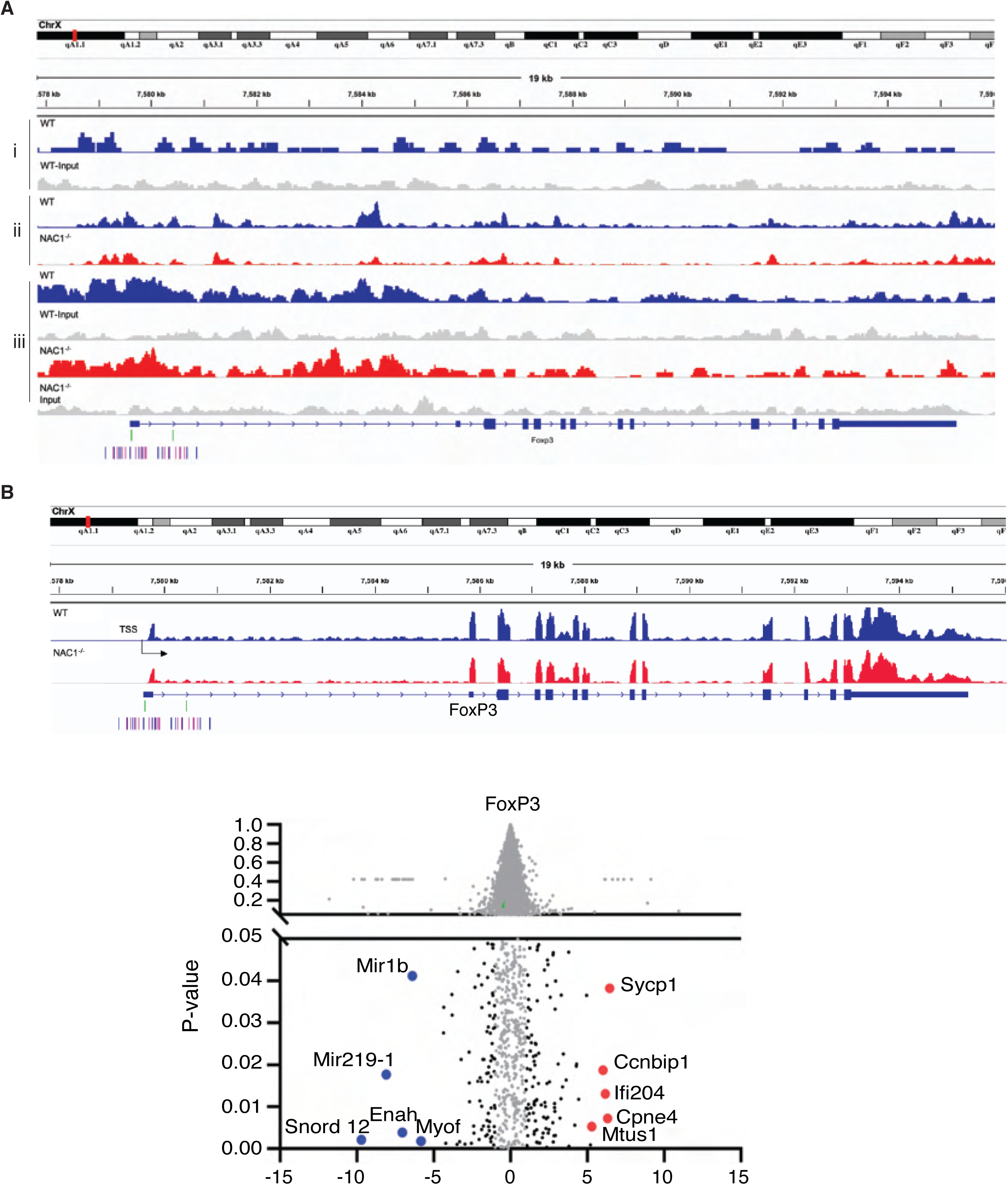
Combined analyses of ChIP-seq, ATAC-seq and RNA-seq indicate that FoxP3 transcription is not regulated by NAC1. (**Ai**) NAC1 ChIP-seq. Approximately 100 differential peaks (NAC1 versus input) were identified using HOMER. NAC1 peak density was not enriched within the regulatory elements of FoxP3. The genes targeted by NAC1-enriched islands in WT Tregs were presented. Representative genomic regions (promotor and CNS1-3) show NAC1 enrichment. Normalized ChIP-seq reads (bigWig) and enriched islands (bed) are shown. (**Aii**) ATAC-Seq. Differential accessibility at FoxP3 gene locus in WT Tregs compared with NAC1^-/-^ Tregs were presented. There were more than 10000 differential ATAC-seq peaks in total, but there were not any significant differential accessibility at FoxP3 gene locus in WT Tregs as compared with NAC1^-/-^ Tregs. (**Aiii**) FoxP3 ChIP-seq. The genes targeted by FoxP3-enriched islands in WT or NAC1^-/-^ Tregs were presented. (**B**) RNA-seq. FoxP3 was not differentially expressed in NAC1^-/-^ Tregs among ∼200 differentially expressed genes, i.e., FoxP3 gene was identically expressed in WT Tregs and NAC1^-/-^ Tregs. All results shown are the representative of three identical experiments.

### NAC1 confines FoxP3 expression by promoting its deacetylation and destabilization

Notably, our comparison of the FoxP3^+^ (YFP^+^) Tregs with the FoxP3^-^ (YFP^-^) CD4^+^ T cells from FoxP3-IRES-mRFP (FIR) reporter mice revealed that the FoxP3^+^ Tregs expressed a low level of NAC1 but a high level of FoxP3; by contrast, the FoxP3^-^ CD4^+^ T cells had a high expression of NAC1 (**Fig. 7A**). Ectopic expression of NAC1 (**Fig. 7B**) in WT Tregs resulted in a reduction of FoxP3 protein (**Fig. 7C**). These results disclose an inverse relationship between the expression of NAC1 and FoxP3 in Tregs. Moreover, co-localization of NAC1 and FoxP3 was observed in the nuclei of Tregs (**Fig. 7D**) and iTregs (**Extended Fig. 7**). These results suggest that NAC1 confines FoxP3 expression.

**Figure 7.**
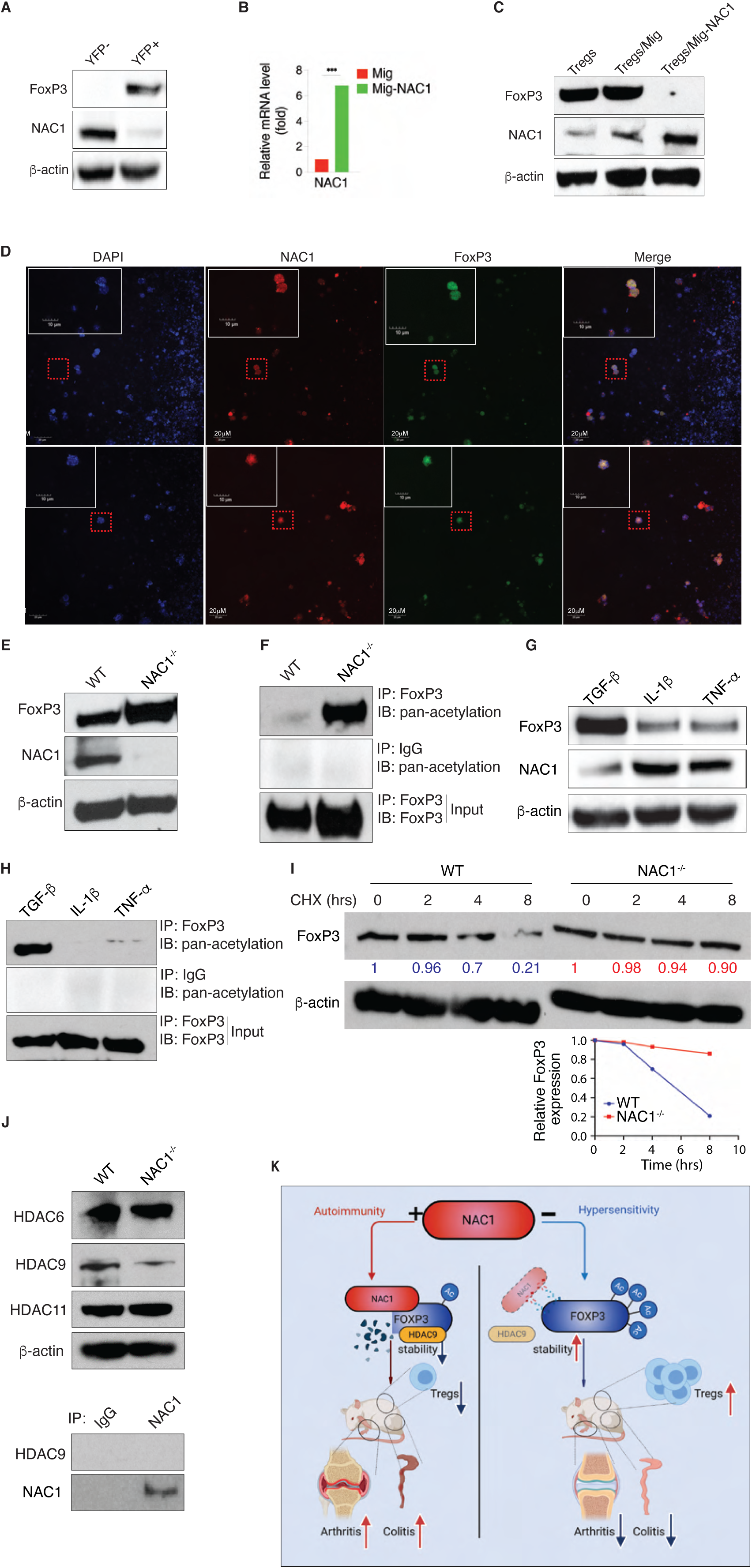
Upregulation of NAC1 by pro-inflammatory cytokines breaks immune tolerance *via* downregulation of FoxP3. (**A**) Expressions of NAC1, FoxP3 and *β*-actin in Tregs from the FIR reporter mice were analyzed by immunoblots. CD4^+^ Tregs from the LNs and spleen of the Foxp3-IRES-mRFP (FIR) reporter mice, WT or NAC1^-/-^ mice, were analyzed. Data shown are the representative of three identical experiments. (**B**) NAC1 mRNA of WT Tregs generated *in vitro* and with ectopic expression of NAC1 were analyzed by RT-PCR. ***, *P*<0.001, Student’s unpaired *t*-test. Data shown are the representative of three identical experiments. (**C**) FoxP3 protein of the sorted Tregs generated *in vitro* and with ectopic expression of NAC1 were analyzed by immunoblots. Data shown are the representative of three identical experiments. (**D**) Immunofluorescent staining of DAPI, NAC1 and FoxP3 in Tregs generated *in vitro*. Data shown are the representative of three identical experiments. **(E)** Expressions of FoxP3, NAC1 and *β*-actin in Tregs from WT and NAC1^-/-^ mice were analyzed by immunoblots. Data shown are the representative of three identical experiments. (**F**) FoxP3 was immunoprecipitated from WT or NAC1^-/-^ Tregs and examined for its Acetylation of Lysine (Pan Acetylation). Acetylated FoxP3 (upper panel), IgG control (middle panel) and FoxP3 (lower panel) were examined by immunoblotting. Data shown are the representative of three identical experiments. (**G**) WT Tregs were cultured in the presence of TGF-*β*, IL-1*β* or TFN-*α* for 12 hours, then the expressions of FoxP3, NAC1 and *β*-actin were analyzed by immunoblots. Data shown are the representative of three identical experiments. (**H**) WT Tregs were cultured in the presence of TGF-*β*1 (2 ng/ml), IL- 1*β* (10 ng/ml) or TFN-*α* (10 ng/ml) for 12 hours, then FoxP3 was immunoprecipitated and examined for its acetylation. Acetylated FoxP3 (upper panel), IgG control (middle panel), and FoxP3 (lower panel) were examined by immunoblots. Data shown are the representative of three identical experiments. (**I**) The pulse-chase experiments. WT or NAC1^-/-^ Tregs were treated with cycloheximide (150 μg/ml) for the indicated hours. FoxP3 protein level was analyzed by immunoblotting (upper panel); a plot indicated that with NAC1 deletion, the stability of FoxP3 was lifted in NAC1^-/-^ Tregs compared to the WT group (lower graph). Data shown are the representative of two identical experiments. **(J)** HDAC expression in WT and NAC1 ^-/-^ Tregs. Expression of HDAC6, HDAC9, HDAC11 and *β*-actin in WT and NAC1^-/-^ Tregs was analyzed by immunoblotting (upper panel). FoxP3 was immunoprecipitated and the immunoprecipitates examined for HDAC9 (lower panel). Data shown are the representative of two identical experiments. (**K**) Proposed model of regulation of FoxP3 by NAC1 in immunity.

Post-translational modifications such as acetylation have a critical role in preventing proteasome-mediated degradation of FoxP3 and maintaining its functional activity (*19, 20*), and in our previous study we found that NAC1 can promote deacetylation of certain protein through its interaction with histone deacetylases (*7*). Thus, the acetylation of FoxP3 protein in NAC1^-/-^ Tregs and WT Tregs was examined. **Fig. 7E** and **Fig. 7F** show that both FoxP3 and acetylated FoxP3 proteins were upregulated in NAC1^-/-^ Tregs as compared with the WT Tregs; in particular, acetylation of FoxP3 was robustly enhanced in NAC1^-/-^ Tregs than that of the WT Tregs (**Fig. 7F**). Conspicuously, in the Tregs treated with the pro-inflammatory cytokines IL-1*β* or TNF-*α*, a vigorous increase of NAC1 expression but evident decreases of both FoxP3 (**Fig. 7G**) and acetylated FoxP3 were observed (**Fig. 7H**). Therefore, it is likely that stabilization of the acetylated FoxP3 protein accounts for the elevated amount of this key transcription factor in NAC1^-/-^ Tregs, and in the Tregs treated with pro-inflammatory cytokines, up-regulation of NAC1 may induce deacetylation and destabilization of FoxP3. Indeed, the pulse-chase experiments demonstrated that the turnover of FoxP3 protein was 4-fold faster in WT Tregs than that in the NAC1^-/-^ Tregs, with a half-life (t_1/2_) of 4 hrs and 8 hrs, respectively (**Fig. 7I).** Histone deacetylases (HDACs) have been reported to modulate the suppressive function of Tregs (*21*); therefore, we determined which HDAC may be involved in the NAC1-induced FoxP3 acetylation. We examined the expressions of HDAC6, HDAC9 and HDAC11, the deacetylases known to affect the stability of FoxP3 and Treg fitness (*22*). We demonstrated the comprised HDAC9 expression in NAC1^-/-^ Tregs as compared to WT Tregs, but did not detect the direct interaction between NAC1 and HDAC9 (**Fig. 7J**). Our results suggest a role of HDAC9 in regulation of the NAC1-mediated FoxP3 stability. Collectively, these results imply that NAC1 negatively regulates FoxP3 stability *via* its effect on deacetylation of this protein, thus weakening immune tolerance.

## Discussion

Autoimmune diseases such as type 1 diabetes, rheumatoid arthritis, ulcerative colitis and Crohn’s disease are presumed to result from interaction between genetic and environmental factors and to be a consequence of compromised immune tolerance versus adaptive immune response; yet, how impaired balance between immune response and tolerance is triggered as well as the mechanisms by which tolerance is established and maintained remain elusive. How the stability of FoxP3 and suppressor Tregs are regulated is an important theme in Treg biology. Using CRISPR screening, a recent study revealed several modulators of FoxP3 expression, and these modulators might be further explored as potential targets for immunotherapy (*23*). In this study, we identified NAC1 as a critical determinant of immune tolerance. We show that NAC1^-/-^ mice is substantial tolerant to the induction of autoimmunity, as evidenced by the significantly decreased occurrences of autoimmune arthritis and colitis (**Fig. 4**). We further show that the promotive effect of NAC1 on autoimmunity is mediated through its negative regulation of the stability of Treg and FoxP3 (**Fig. 7**).

Although DNA methylation of FoxP3 has been reported to be associated with the stability and function of Tregs (*24–27*), the effects of NAC1 on Tregs do not appear to be associated with alterations in DNA methylation of FoxP3. We compared the DNA methylation of FoxP3 in WT and NAC1^-/-^ Tregs and examined the DNA methylation of FoxP3 in the Treg-specific demethylated region (TSDR) of CNS2 (ADS443) and FoxP3 proximal promoter region (ADS1183) (**Fig. 5**). As a control, FoxP3 DNA methylation was similar in the unsorted lymphocytes from the LNs and spleen of NAC1^-/-^ mice (**Extended Data Table 1**). In these experiments, greater than 100 CpG sites in the FoxP3 DNA promoter regions showed that there was no significant difference between WT and NAC1^-/-^ Tregs, indicating that NAC1 does not affect the DNA methylation of FoxP3. Moreover, we found that NAC1 does not act as a transcriptional regulator (**Fig. 6**). Instead, we show that NAC1 can interfere the acetylation and degradation of FoxP3 protein. (**Fig. 7**). Thus, the role of NAC1 in the post-transcriptional regulation of FoxP3 may account for the upregulation of FoxP3 in NAC1^-/-^ Tregs.

Treg stability is vital to the maintenance of immune tolerance but is often altered in autoimmunity; yet, how destabilization of Tregs occurs in autoimmune diseases remains elusive. In autoimmune arthritis, TNF-*α* plays a more important role in triggering events leading to inflammation both locally and systemically, whereas IL-1 is more involved at the local level in the processes leading to cartilage and bone destruction and in impeding cartilage repair. Nevertheless, IL-1 and TNF-*α* strongly synergize in numerous biological functions, and simultaneous blockade of IL-1 and TNF-*α* provides favorable effects in suppressing arthritis development, suggesting the importance of both of the cytokines (*28, 29*). Our results reported here, which show the concomitant upregulation of NAC1 and downregulation of FoxP3 in Tregs treated with the pro-inflammatory cytokines such as IL-1*β* and TNF-*α* (**Fig. 7H**), provide at least a partial explanation for this question. Based on these findings, we speculate that the “basal” level of NAC1 in Tregs plays an important role in leashing the immune tolerance to keep the immune system vigilant to pathogens; inflammatory stimulation induces upregulation of NAC1, and this in turn destabilizes FoxP3 and converts FoxP3^+^ Tregs to FoxP3^-^ Tregs that then become Th1 or Th17 CD4^+^ Teffs, further breaking tolerance and instigating strong immune response (**Fig. 7K**).

In this study, the role and importance of the NAC1-mediated regulation of FoxP3 in Tregs were largely determined in mice with complete knockout of NAC1 gene. We demonstrated that NAC1 can modulate the stability of FoxP3 protein, and loss of NAC1 enhances the functional activity of Tregs. It shall be interesting to further investigate the functions of NAC1 in hematopoietic cells. Our data showing that NAC1 deletion is associated with reduced effector cell activity (**Extended Data Figs. 2 and 3)** open a question of whether impairment of the effector responses is also involved in the reduced inflammation caused by NAC1 deletion. The roles of NAC1 in other immune cells such as CD8^+^ T and CD4^+^ Th1 or Th17 effector cells warrant further investigation.

Because the mice that we used in this study were subjected to complete knockout of NAC1 gene, the deficiency of NAC1 in other cell types may also affect immune tolerance. For example, the role of NAC1 in other types of T cells such as CD8^+^ T cells, conventional CD4^+^ T cells (i.e., Th1, Th2, Th17) and innate immune cells including macrophages and dendritic cells remain unclear. To exclude the possibility that NAC1 deficiency in other types of cells may impact Treg development, we performed the bone marrow chimera experiment in which the bone marrow cells (CD4^-^CD8^-^) from WT and NAC1^-/-^ mice were transferred into X-ray-irradiated WT mice, and 6 weeks later, we euthanized the mice and isolated the spleen, LNs and thymus for examining Treg development. These experiments showed the comparable numbers of Tregs in the thymus of the chimera receiving either WT or NAC1^-/-^ bone marrow cells. However, we observed that the bone marrow transplants from NAC1^-/-^ mice generated greater numbers of Tregs in the LNs and spleen than those from WT (**Extended Fig. 8**), which is similar to that we classified in NAC1^-/-^ mice. This result also supports the conclusion that in the peripheral system, NAC1^-/-^ Tregs are more stable than WT cells.

As the stability of Tregs is vital to maintenance of immune tolerance, the role of NAC1 in destabilization of suppressor Tregs provides a promising opportunity for therapeutic manipulation of their stability and function. We believe that therapeutic targeting of NAC1 to revitalize suppressor Tregs may be further exploited as a potentially novel tolerogenic strategy to treating autoimmune diseases. We have recently identified and characterized a small molecule inhibitor of NAC1, NIC3, through a high-throughput screening, and showed that this compound can effectively promote the proteasome-mediated degradation of NAC1 protein (*30*). We anticipate that inhibiting NAC1 by pharmacologic approaches shall provide further insights into the feasibility and effectiveness of NAC1-based modulation of Tregs stability for therapeutic purposes.

## Methods

### Cell lines and mice

C57BL/6 (B6), *Rag1^-/-^* and FOXP3-IRES-mRFP (FIR) reporter mice were purchased from The Jackson Laboratory (Bar Harbor, ME). NAC1^-/-^ mice were generated by Dr. Jian-long Wang and crossed in the C57BL/6 background for more 10 generations (*6, 31*). All the animal experiments were performed in compliance with the regulations of The Texas A&M University Animal Care Committee (IACUC) and in accordance with the guidelines of the Association for the Assessment and Accreditation of Laboratory Animal Care.

### T cell culture

T cells were cultured in 48-well plates containing 1 ml RPMI 1640 (Invitrogen) with 10% fetal calf serum (Omega Scientific, CA). T cell isolation kits including mouse CD4^+^ (# 130-104-454), CD8a^+^ (#130-104-075) and CD4^+^ CD25^+^ Treg (#130-091-041), T cell activation/expansion kit (#130-093-627) and Treg expansion kit (#130-095-925) were purchased from the Miltenyi Biotec (Auburn, CA). Recombinant mouse TGF-*β* (#763104), IL-1*β* (#575106), and TNF-*α* (#575206) were obtained from BioLegend (San Diego, CA).

### Cytokine secretion, cell recovery, and proliferation/cell division

IL-2 and IFN-*γ* were measured using ELISA after 48 hr of culture (*32*). Latent TGF-*β*1 (#433007, BioLegend) and IL-10 (#431411, BioLegend) were determined using ELISA after 48 hr of stimulation. *In vitro* T cell survival was determined using trypan blue exclusion. Proliferation/division of T cells was measured using the CellTrace™ CFSE Cell Proliferation Kit (#C34554, Invitrogen).

### Metabolic assays

Purified CD4^+^ Tregs were plated in the Cell-Tak-coated Seahorse Bioanalyzer XFe96 culture plates (300,000 or 100,000 cells/well, respectively) in assay medium consisting of minimal, unbuffered DMEM supplemented with 1% BSA and 25 mM glucose, 2 mM glutamine (and 1 mM sodium pyruvate for some experiments). Basal rates were taken for 30 min, and then streptavidin- complexed anti-CD3bio at 3 mg/mL ± anti-CD28 at 2 mg/mL or PMA (CAS 16561-29-8) (Fisher) was injected and readings were taken for 1–6 hr. In some experiments, oligomycin (2 mM), carbonyl cyanide p-trifluoromethoxyphenylhydrazone (FCCP) (0.5 mM), 2-deoxy-d-glucose (10 mM) and rotenone/antimycin A (0.5 mM) were injected to obtain maximal respiratory and control values. Because ECAR values tend to vary among experiments, both a representative trace and normalized data (calculated as the difference between maximal and basal ECAR values) were shown in the figures.

### In vitro mouse Treg generation

Naive CD4^+^CD25^-^ T cells from the LNs and spleen of WT or NAC1^-/-^ mice were incubated with the indicated reagents including TGF-*β* in the CellXVivo™ Mouse Treg Cell Differentiation Kit (#CDK007, R&D Systems) for 5 days.

### In vitro Treg suppression assay

CD4^+^ CD25^+^ Tregs were co-cultured with the CFSE-labeling CD4^+^ CD25^-^ responder T cells from the pooled LNs and spleen of C57BL/6 mice in various ratios. To stimulate T cells, the mixed T cells were treated with the T cell activation/expansion kit (#130-093-627; Miltenyi Biotec). As controls, CD4^+^ CD25^+^ Tregs and CD4^+^ CD25^-^ responder T cells were cultured without any stimulus. Suppression of responder T cells was determined by measuring CFSE dilution.

### T cell transfer model of colitis

Naive CD4^+^ T effectors (Teffs, CD45RB^hi^CD25^−^) from B6 mice and CD4^+^ Tregs (CD45RB^lo^CD25^+^) from WT or NAC1^-/-^ mice were purified using a high-speed cell sorter. Naive CD4^+^ Teffs (6× 10^5^ cells/mouse) without or with Tregs (2× 10^5^ cells/mouse) were then *i.p.* transferred into *Rag*1^-/-^ mice. Body weights were recorded twice a week. When loss of body weight exceeded 20% after transfer, the host mice were sacrificed.

### Retroviral transduction

Full-length cDNA of NAC1 was provided by Dr. Ie-Ming Shih and Tian-Li Wang (John Hopkins University (*33*), and subcloned into the Mig vector containing GFP for retroviral transduction of mouse Tregs (*34*).

### Antibodies and reagents

PE-, PE/Cy7, Alexa 647, APC or APC/Cy7-conjugated anti-mouse CD4 (GK1.5), CD8 (53-6.7), CD25 (3C7), CD45RB (C363-16A), CD25 (3C7), CD44 (IM7), CD117 (2B8), TCRV*β* (H57-597), TGF-*β*1(TW7-16B4), FoxP3 (MF-14) and Acetylated Lysine (15G10; #623402) were purchased from BioLegend (San Diego, CA). Rabbit NAC1 (#4183), HDAC6 (#7612) and *β*-actin (#8457) antibodies were purchased from Cell Signaling (Beverly, MA). Rabbit anti-NAC1 antibody (ab29047) for immunoprecipitation was obtained from Abcam (Cambridge, MA). Mouse HDAC9 (#sc-398003) and HDAC11 (#sc-390737) antibodies were purchased from Santa Cruz Biotech (Dallas, TX). Cycloheximide were purchased from Sigma-Aldrich Corporation (Sigma-Aldrich, St Louis, MI).

#### Immunoprecipitation and Immunoblotting

Cells were lysed in ice-cold RIPA Lysis Buffer (#89900, Thermo Scientific, MA) for 30 min. Insoluble materials were removed, and the lysates were used for Western blotting or immunoprecipitated overnight with an antibody such as anti-FoxP3 antibody followed by incubation with protein G agarose beads at 4°C for 2 hr. The washed immunoprecipitates were boiled in SDS sample buffer and the protein content was determined by Bio-Rad protein assay kit (#5000002, Bio-Rad, Hercules, CA). Equal amounts (30-50 µg) were loaded onto 4-12% NuPage Bis-Tris pre-casting gels (SDS-PAGE), transferred onto PVDF membrane (Invitrogen), and immunoblotted. All blots were developed with the ECL immunodetection system (#426319, BioLegend, CA).

### RT-PCR

Retrovirally transduced Tregs with Mig or Mig-NAC1 were unsorted or sorted, and total RNA was extracted from the Tregs using QIAgen RNeasy mini kits. Samples were subjected to reverse transcription using a high-capacity cDNA synthesis kit (Applied Biosystems). PCR analysis was performed using TaqMan real-time PCR (Thermo Fisher Scientific). Primers used are: FoxP3 forward: 5’-CCCAGGAAAGACAGCAACCTT-3’, FoxP3 reverse: 5’-TTCTCACAACCAGGCCACTTG-3’; NAC1 forward: 5’-TGC TTA GTT AAC TTA CTG CAG GGC TTC AGC CGA-3’, NAC1 reverse: 5’-TAA GCA CTC GAG ATG GCC CAG ACA CTG CAG ATG-3’.

### CpG DNA methylation

CpG DNA methylation was analyzed by bisulphite treatment of RNase-treated genomic DNA, followed by PCR amplification and pyrosequencing (Pyro Q-CpG), which was performed by EpigenDX. Eight mouse genes, including FoxP3, Ctla4, Ikzf2, Ikzf4, Tnfrsf18, Il2ra, Cd274 and Irf4, were screened for methylation percentage in various regulatory regions. Sequences analyses for FoxP3 were: *FoxP3* promoter, *FoxP3* CNS2 and *FoxP3* 3′ region.

### RNA-Seq

Tregs were mechanically disrupted and homogenized using a Mini-BeadBeater-8 (BioSpec Products, Bartlesville, Oklahoma). RNA was extracted using a RNeasy Mini Kit (Qiagen, Valencia, California). RNA concentration and integrity were measured using an Agilent 2100 Bioanalyzer (Agilent Technologies, Santa Clara, CA). All samples had an RNA Integrity Value (RIN) of > 7.5. RNA-Seq libraries were prepared using the Illumina TruSeq Stranded mRNA Library Prep Kit (Illumina, San Diego, CA) and sequenced on an Illumina HiSeq 2500 Sequencer (Illumina, San Diego, CA) as 75 base pair (bp) paired-end reads.

### CHIP-seq

ChIP was performed as described (*35*), with some modifications. Tregs were subjected to sonication using a Bioruptor® Pico sonication device (Diagenode) to obtain 100–500-bp chromatin fragments. A total of 250 µg of sonicated chromatin fragments were incubated with 10 µg of NAC1 antibody for crosslinking with magnetic beads (no. 11201D, Dynabeads® M280 sheep anti-mouse IgG, Dynal Biotech, Invitrogen). The cross-linked samples were reversed at 65°C for overnight, and the precipitated DNA was treated with RNase A and proteinase K, and then purified using the QIAquick PCR purification kit (QIAGEN). The DNA libraries were prepared following the guidelines from Illumina (Fasteris Life Sciences; Plan-les-Ouates, Switzerland). Input DNA was sequenced and used as a control. The DNA libraries were sequenced on Illumina HiSeq2500, producing 25–35 million reads per sample.

### ATAC-Seq

Tregs were freshly dissected and processed for ATAC-seq. In brief, the tissues were resuspended in 1 ml of lysis buffer (1× PBS, 0.2% NP-40, 5% BSA, 1 mM DTT, protease inhibitors), followed by Dounce homogenization with a loose pestle using 20 strokes. The lysates were then filtered through a 40-μm cell strainer, and the nuclei were collected by centrifugation at 500*g* for 5 min. Tagmentation was performed immediately according to the reported ATAC-seq protocol (*36*).

### Pulse-chase analysis

Isolated Tregs from WT or NAC1^-/-^ mice were activated and expanded with kits (# 130-104-454 and #130-095-925; Miltenyi Biotec), then treated with cycloheximide (150 μg/ml) for various periods of time. FoxP3 protein was analyzed by immunoblotting.

### Collagen-induced arthritis

C57BL/6 mice (4 months old) were injected at the base of the tail with 0.1 mL of emulsion containing 100 µg of bovine type II collagen (CII) (Chondrex, Redmond, WA, USA) in complete Freund’s adjuvant (CFA) (Chondrex), using a 1-mL glass tuberculin syringe with a 26-guage needle. Mice were assessed for arthritis in the paws (*37*).

### DSS-induced colitis

Colitis was induced in mice by oral ingestion of 3% dextran sulfate sodium (DSS, SKU 02160110-CF; MP Biomedicals) in drinking water for 5 days. The severity of colitis activity was graded on designated dates as described (*38*). Body weight, occult or gross rectal bleeding, and feces consistency (on scales of 0–4) were monitored for each mouse. The resultant IBD disease activity index is the average of the scores of the colitis symptoms. The occult blood in mouse fecal samples was detected using Hemoccult Test Kit (Beckman Coulter Inc, Fullerton, CA).

### Histology and immunohistochemistry

Joint or gut tissues were fixed with 10% neutral formalin solution (VWR, West Chester, PA), and the fixed samples were prepared and stained with H&E as described (*39*). For immunofluorescent microscopy, the tissues were frozen in cryovials on dry ice immediately following resection. Cryo- sectioning and immunofluorescent staining were performed as described (*39*).

### Statistical analysis

Multiple unpaired *t*-test, simple linear regression and survival curve comparison were performed to analyze the differences between the groups, using GraphPad Prism (GraphPad Software, San Diego, CA); significance was set at 5%.

## Extended Methods

### Reagents

H-2K^b^ VACV B8R (TSYKFESV) Tetramer (#TB-M538-1, MBL), anti-mouse CD36 antibody (clone HM36, BioLegend), anti-human/mouse/rat NAC1 antibody (clone SWN-3, BioLegend), anti-human FoxP3 antibody (clone 206D, BioLegend), and L-(+)-Lactic acid (#ICN19022805, MP Biomedicals).

### Viral infection

VACV infection was performed by an intraperitoneal injection of viruses (2×10^6^ PFU/mouse) as described (*40*).

### Murine Melanoma Model

WT or NAC1^-/-^ Tregs (1 x 10^5^) were injected *s.c.*in the flank region of the recipient mice inoculated with 1 x 10^6^ B16 tumor cells. Tumor sizes were measured by a caliper and tumor volumes were calculated as: V=long diameter × short diameter^2^ × 0.52 (*41*).

#### LPS-induced colitis

To induce colitis, B6.Thy1.1 Tg mice were *i.p.* injected with 20 μg/kg LPS. In the following day, 3x10^6^ WT or NAC1^-/-^ Tregs were *i.v.* transferred into LPS-challenged mice *via* the tail vein. Seven days later, mice were sacrificed and the colons and spleens of the mice were dissociated. The length of colon was measured, and the colons were processed for H&E staining and flow cytometric analysis.

#### Generation of bone marrow chimera

To prepare B6. Thy1.1^+^ Tg recipient mice for irradiation, drinking water was removed 12 hrs prior to irradiation. The mice were then X-ray irradiated (1000 rad) with a RS200 X-ray irradiator (Rad Source Technologies Inc. GA, USA). The bone marrow cells were harvested from femur of C57BL/6 Thy1.2^+^ WT and NAC1^-/-^ mice, using standard procedures. Red blood cells (RBCs) were lysed, and CD4^+^ cells and CD8^+^ cells were depleted by the negative selection using biotin conjugated anti-CD4 and anti-CD8 antibodies and streptavidin nanobeads. CD4^+^ and CD8^+^- depletions were analyzed by flow cytometry. Ten millions of CD4/CD8-depleted WT and NAC1^-/-^ bone marrow cells were *i.v.* injected into the irradiated B6.Thy1.1^+^ recipient mice. Sulfatrim (sulfamethoxazole and trimethoprim oral suspension)-containing drinking water (5ml/200ml) was given to the irradiated animals for 2 weeks. After 6 weeks, the spleen, LNs and thymus were isolated, and cells were analyzed by flow cytometry to evaluate the development of Tregs.

## Author contributions

J.Y. and J.S. designed the experiments, analyzed the data, and wrote this paper. Y.R., A.K., X.X., JK.D., H.P., L.W., Y.Z. C.J., Y.C., and L.Z. performed the experiments. D.F., RC.A., P.dF. and J. W. provided reagents or NAC1^-/-^ mice. X.X., JK.D., H.P., Y.Z., C.J., L.Z., X.L and Y.C. analyzed the data.

## Acknowledgments

This work was supported by National Cancer Institute Grant R01CA221867 to JM. Yang and J.S., and National Institute of Health Grant R01AI121180 and R21AI128325, the American Diabetes Association (1-16-IBS-281), and Department of Defense Grant LC210150 to J.S.

## Guarantor’s statement

J.Y. and J.S. are the guarantors of this work and, as such, had full access to all the data in the study and takes responsibility for the integrity of the data and the accuracy of the data analysis.

## Legends to Extended Data

**Extended Data Figure 1.**
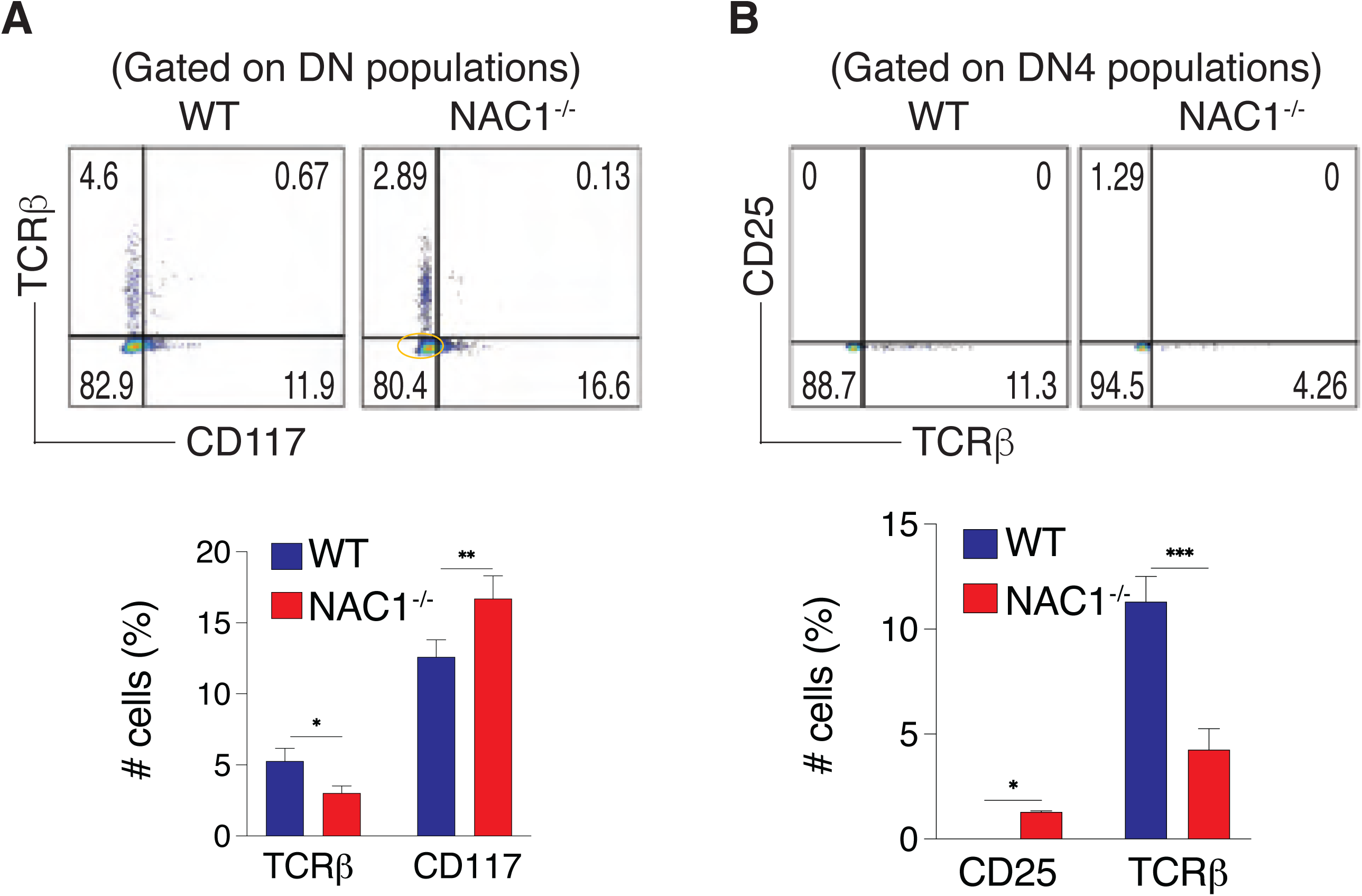
NAC1^-/-^ mice have a decreased percentage of TCRVβ cells in DN4 stage. The thymocytes from WT or NAC1^-/-^ mice were analyzed by flow cytometry and calculated for numbers or percentages. (**A**) CD117 and TCR V*β*. The DN populations were analyzed for CD117 and TCRV*β*. Data shown are the representative of three identical experiments. (**B**) TCRV*β* and CD25. The DN4 populations were analyzed for TCR V*β* and CD25. Data are the representative of three identical experiments (N = 5). **\*,** *p*<0.05, **, *p*<0.01, ***, *p*<0.001, Student’s unpaired *t*- test.

**Extended Data Figure 2.**
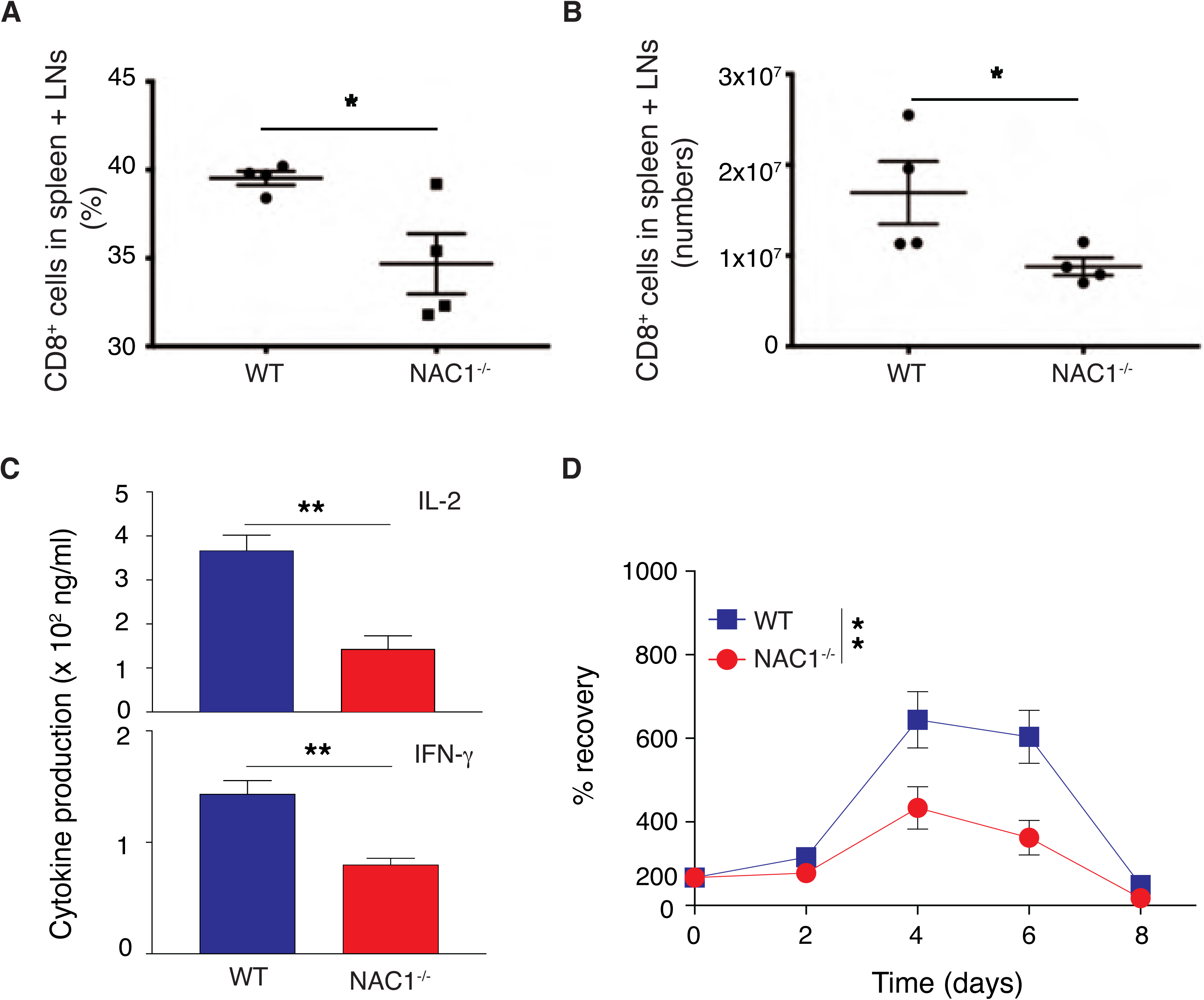
NAC1^-/-^ CD8^+^ T cells are defective in cytokine production and survival. (**A-B**) Percentages (**A**) and numbers (**B**) of CD8^+^ T cells from the pooled LNs and spleen of WT or NAC1^-/-^ mice. Data shown are the representative of three identical experiments. The values represent mean ± S.D. (N = 3). *, *p*<0.05, Student’s unpaired *t*-test. (**C-D**) Purified CD8^+^ T cells from the pooled LNs and spleen of WT or NAC1^-/-^ mice were stimulated with anti- CD3 plus CD28 antibodies. (**C**) Cytokine production. ** *p*<0.01, Student’s unpaired *t*-test. The values represent the mean ± S.D. (N = 3). (**D**) Cell recovery on various days. The numbers of T cells present on day 0 were assigned a value of 100%, and numbers surviving on various days were used to calculate the percentage recovery relative to day 0. Data shown are the mean ± S.E.M. of percentage change of a representative of three identical experiments (N = 3). **, *p*<0.01, Nested *t*-test).

**Extended Data Figure 3.**
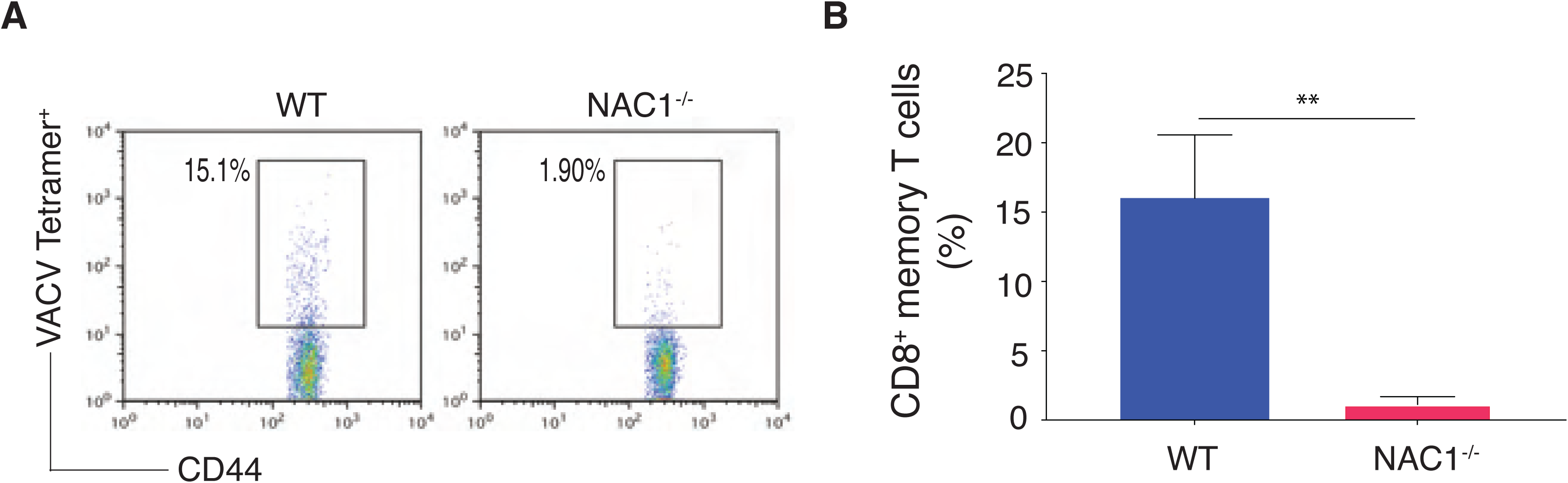
Memory CD8^+^ T cell development is altered in NAC1^-/-^ mice. WT or NAC1^-/-^ mice were challenged with VACV (*i.p.*) for 35 days, then the pooled LNs and spleen were analyzed by flow cytometry. (**A**) CD44 and VACV tetramer staining, gating on CD8^+^CD44^+^ population. (**B**) Quantification of percentage of memory CD8^+^ T cell populations. The results shown are the mean ± S.E.M. of three identical experiments (N = 5). ** *P*<0.01, Student’s unpaired *t*-test.

**Extended Data Figure 4.**
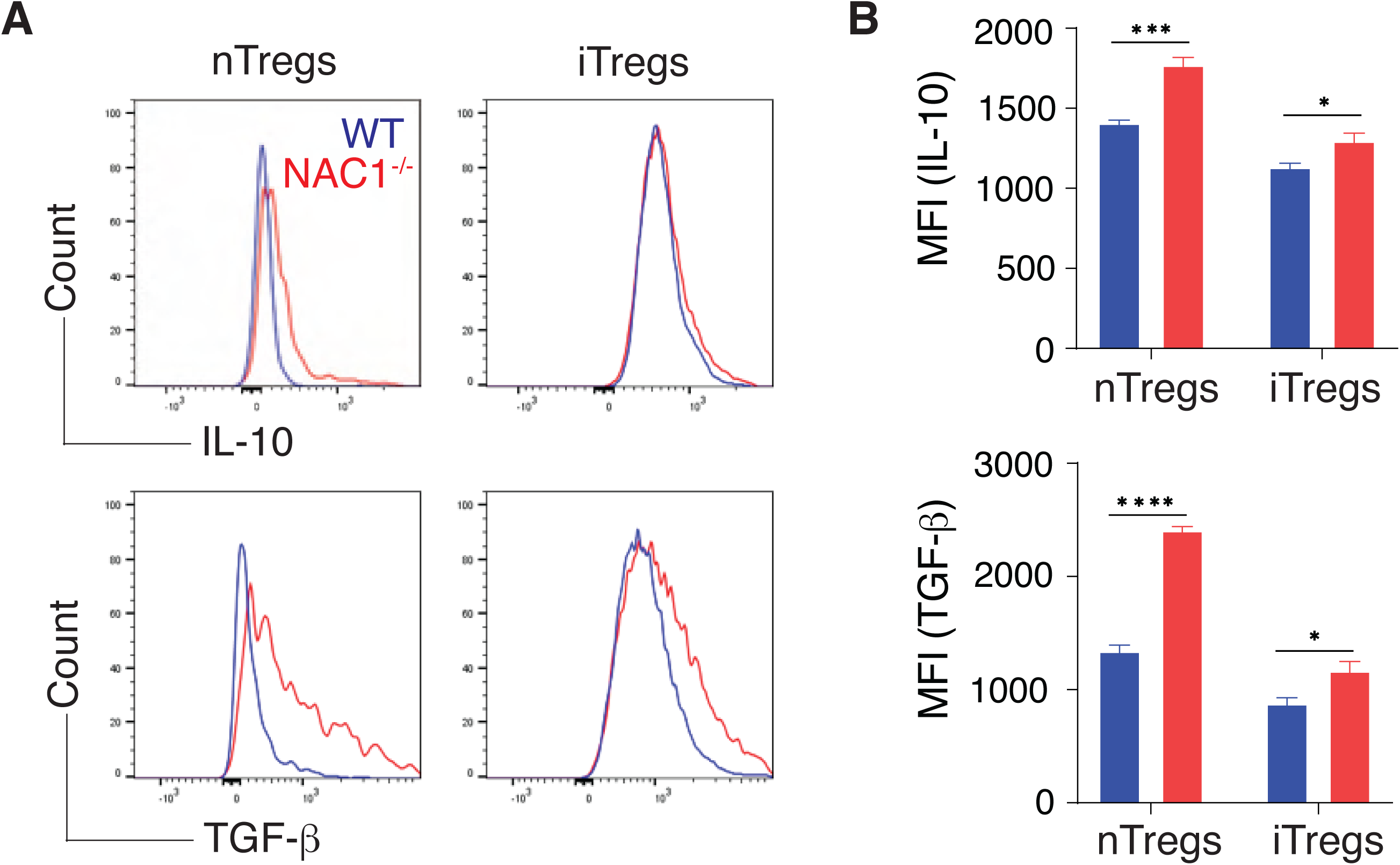
Both NAC1^-/-^ iTregs and nTregs produce more suppressive cytokines than WT iTregs and nTregs. WT or NAC1^-/-^ iTregs and nTregs were examined productions of suppressive cytokines (IL-10 and TGF-*β*) by intracellular staining and flow cytometric analysis. (**A**) Productions of IL-10 and TGF-*β*. (**B**) MFI of IL-10 and TGF-*β*. Data shown are the representative of three identical experiments. * *P*<0.05, ***, *P*<0.0001, ***, *P*<0.00001, Student’s unpaired *t*-test.

**Extended Data Figure 5.**
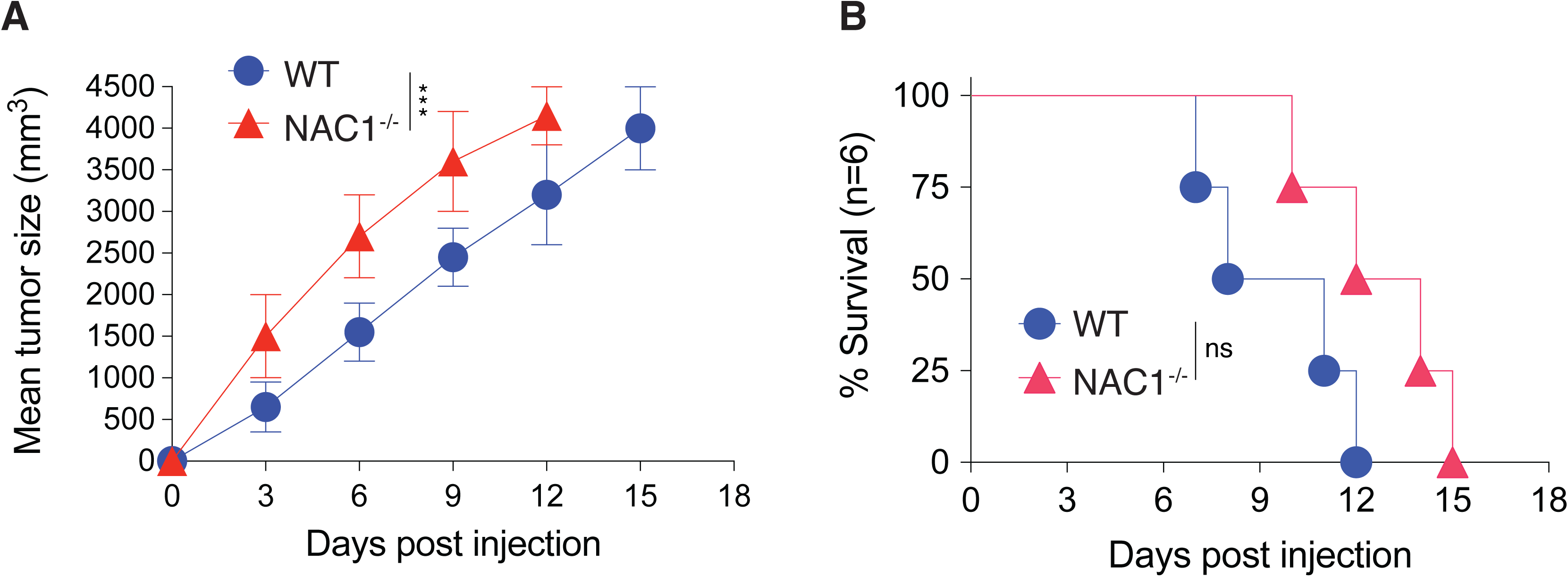
NAC1^-/-^ Tregs show enhanced suppressive function. WT or NAC1^-/-^ Tregs (1 x 10^5^) were injected *s.c.*in the flank region of the recipient mice with 1 x 10^6^ B16 tumor cells on various days. (**A**) Tumor growth. Data shown are the mean ± S.E.M. of tumor sizes of a representative of three identical experiments (N = 6). ***, *P*<0.0001, simple linear regression. (**B**) Survival curves. Data shown are the representative of three identical experiments (N = 6). ns, *P*>0.05, Log-rank (Mantel-Cox) test.

**Extended Data Figure 6.**
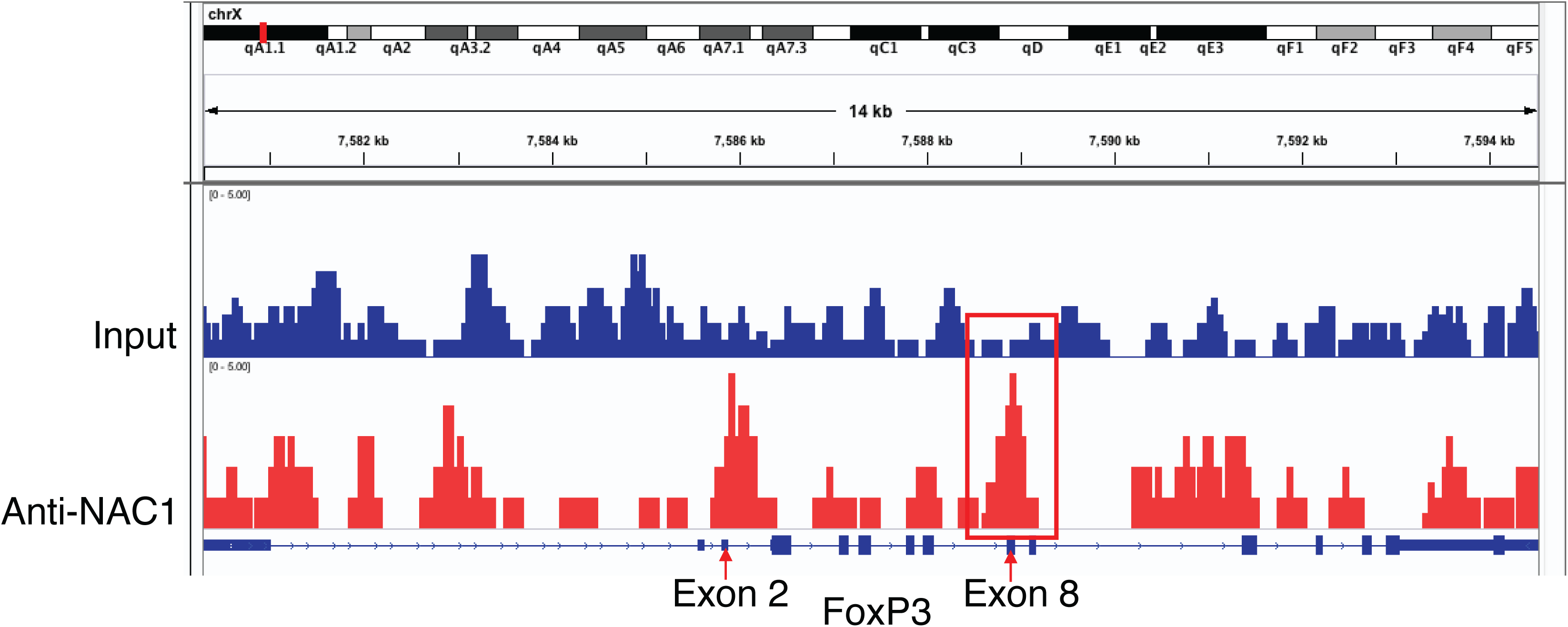
FoxP3 elements regulated by NAC1 are located between Exon 2 and Exon 8. ChIP-seq analysis of naive CD4^+^CD25^+^ Tregs from the pooled LNs and spleen of WT mice. NAC1-enriched islands are shown. Representative genomic regions (Exon 2 and Exon 8) show NAC1 enrichment. Normalized ChIP-seq reads (bigWig) and enriched islands (bed) are shown. Results shown are the representative of three identical experiments

**Extended Data Figure 7.**
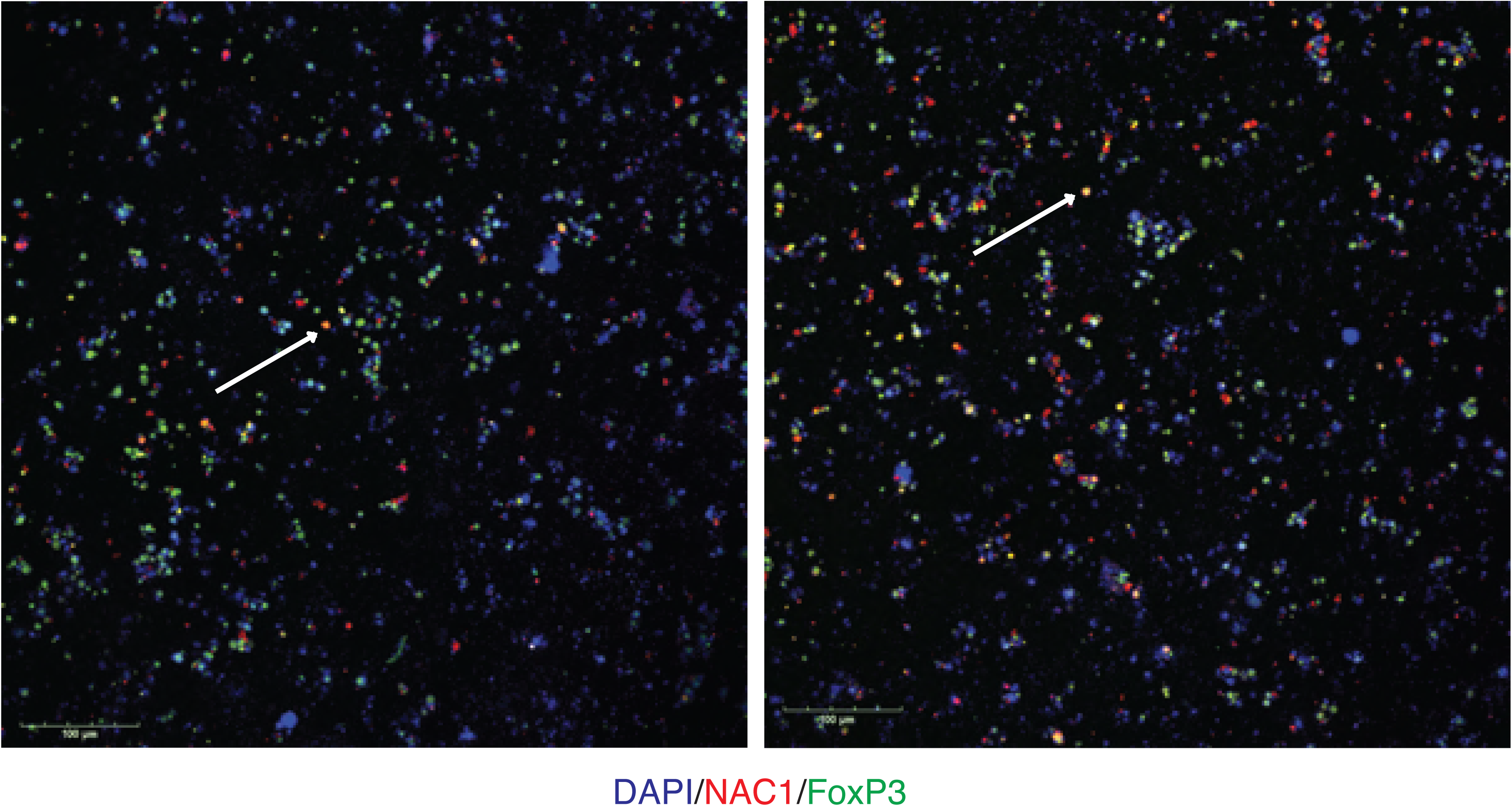
Co-localization of NAC1 and FoxP3 in the nuclei of Tregs. Immunofluorescent staining of DAPI, NAC1 and FoxP3 in Tregs generated *in vitro*. Arrows indicate overlays with DAPI, NAC1 and FoxP3. Data shown are the representatives of three identical experiments.

**Extended Data Figure 8.**
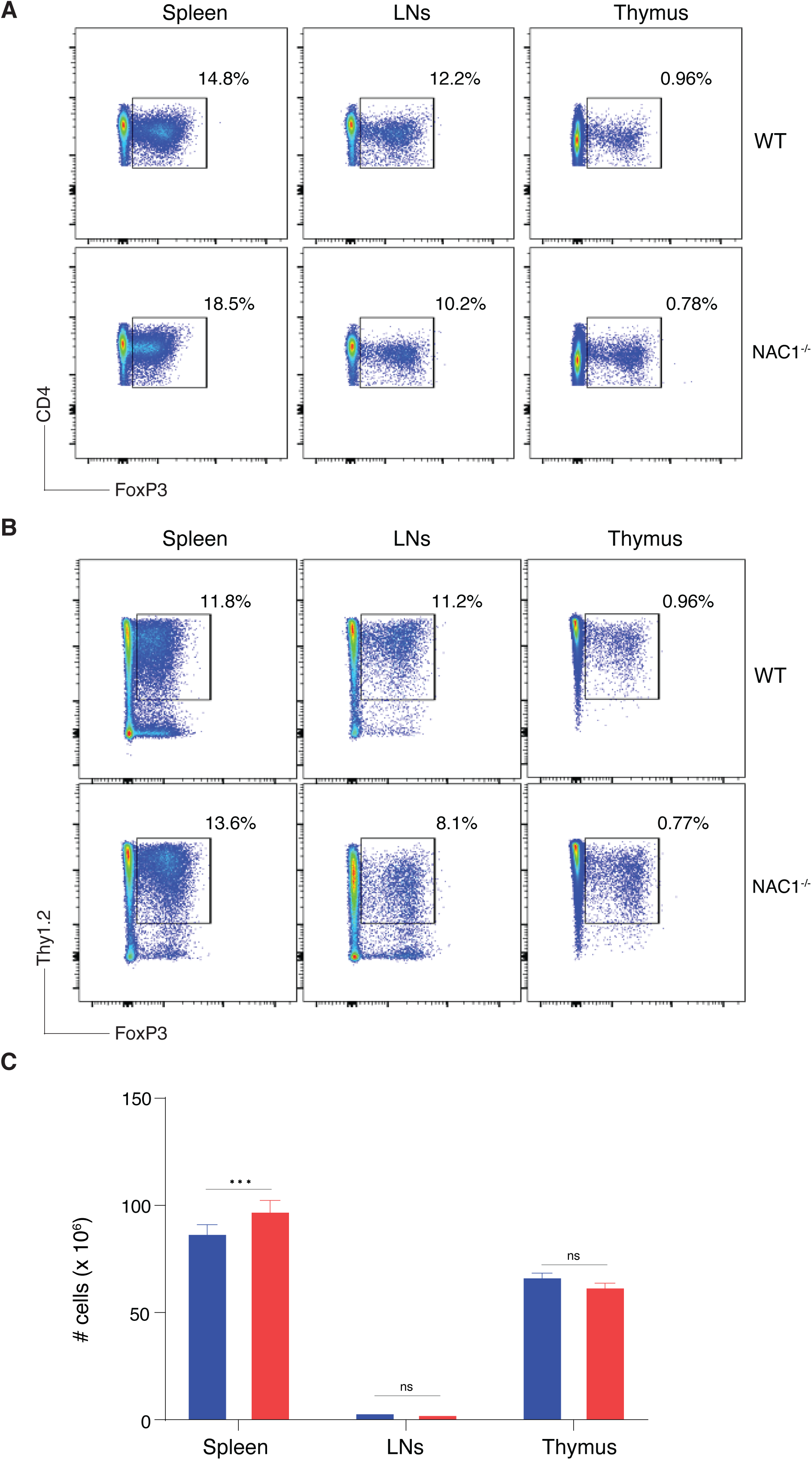
Thymic development of NAC1^-/-^ Tregs in the bone marrow chimeras. Bone marrow cells (CD4^-^CD8^-^; Thy1.2^+^) from WT and NAC1^-/-^ mice were transferred into X-ray irradiated mice (Thy1.1^+^). Six weeks later, the mice were euthanized, and the spleen, LNs and thymus were isolated to examine Treg development using flow cytometry. (**A**) CD4^+^FoxP3^+^ cells. Data shown are the representatives of two identical experiments (N = 10). (**B**) Thy1.1^+^FoxP3^+^ cells. Data shown are the representatives of two identical experiments (N = 10). (**C**) Total numbers of cells. Data shown are the mean ± S.E.M. of a representative of two identical experiments (N = 10). ***, *p*<0.001; ns, no statistical difference, Student’s unpaired *t*-test).

**Extended Data Table 1.**
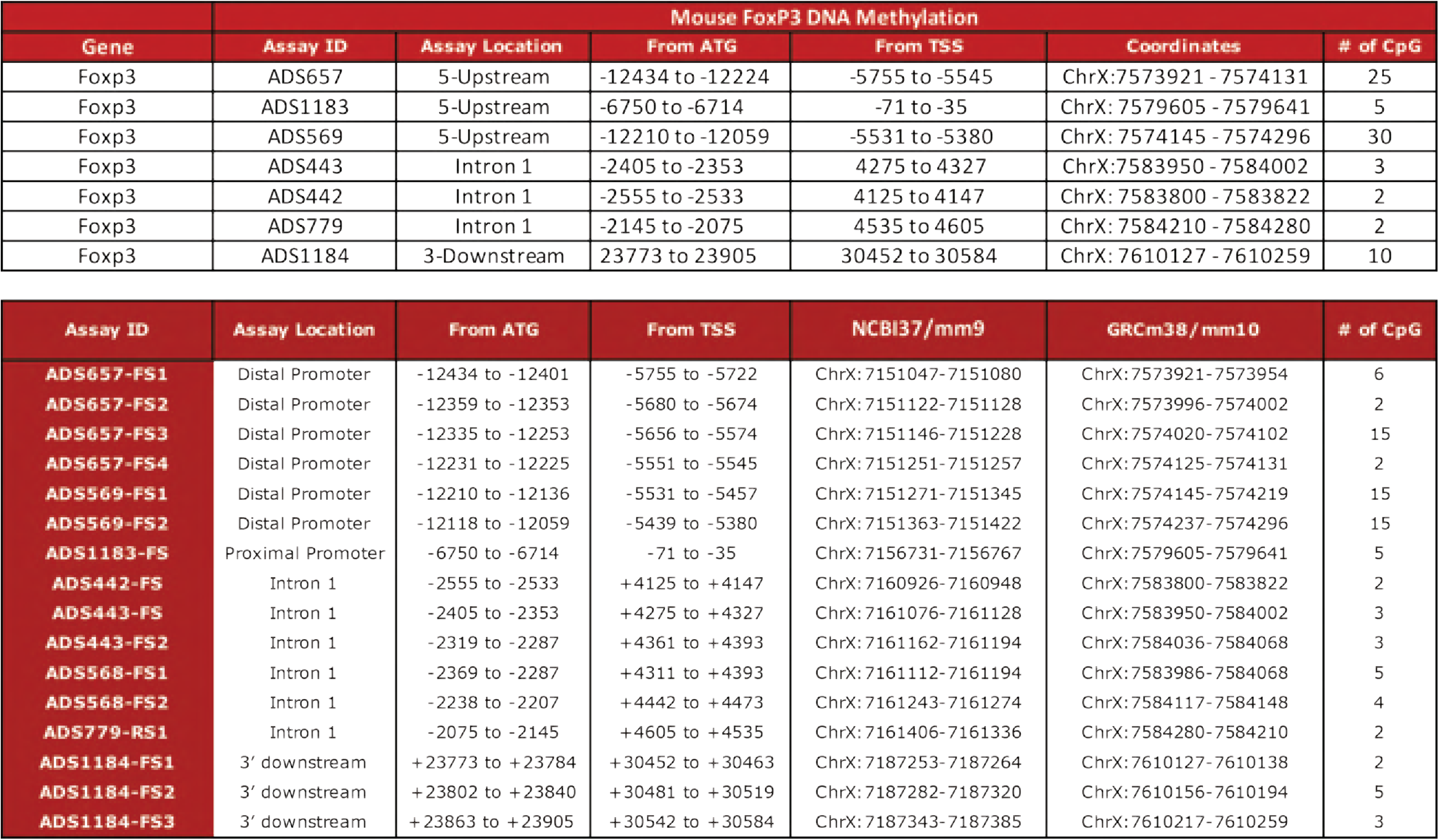
Detailed CpG analysis of mouse FoxP3 DNA methylation. Seven CpG sites of FoxP3 regulators were analyzed.

